# Live-cell fluorescence imaging and optogenetic control of PKA kinase activity in fission yeast *Schizosaccharomyces pombe*

**DOI:** 10.1101/2024.01.14.575615

**Authors:** Keiichiro Sakai, Kazuhiro Aoki, Yuhei Goto

**Affiliations:** Quantitative Biology Research Group, Exploratory Research Center on Life and Living Systems (ExCELLS), National Institutes of Natural Sciences, 5-1 Higashiyama, Myodaiji-cho, Okazaki, Aichi 444-8787, Japan; Division of Quantitative Biology, National Institute for Basic Biology, National Institutes of Natural Sciences, 5-1 Higashiyama, Myodaiji-cho, Okazaki, Aichi 444-8787, Japan; Department of Basic Biology, School of Life Science, SOKENDAI (The Graduate University for Advanced Studies), 5-1 Higashiyama, Myodaiji-cho, Okazaki, Aichi 444-8787, Japan; Center for Living Systems Information Science, Graduate School of Biostudies, Kyoto University, Yoshida-Konoecho, Sakyo-ku, Kyoto, Kyoto 606-8501, Japan; Laboratory of Cell Cycle Regulation, Graduate School of Biostudies, Kyoto University, Yoshida-Konoecho, Sakyo-ku, Kyoto, Kyoto 606-8501, Japan

**Keywords:** fission yeast, cAMP-PKA pathway, kinase translocation reporter (KTR), optogenetics

## Abstract

The cAMP-PKA signaling pathway plays a crucial role in sensing and responding to nutrient availability in the fission yeast *Schizosaccharomyces pombe.* This pathway monitors external glucose levels to control cell growth and sexual differentiation. However, the temporal dynamics of the cAMP-PKA pathway in response to external stimuli remains unclear mainly due to the lack of tools to quantitatively visualize the activity of the pathway. Here, we report the development of the kinase translocation reporter (KTR)-based biosensor spPKA-KTR1.0, which allows us to measure the dynamics of PKA activity in fission yeast cells. The spPKA-KTR1.0 is derived from the transcription factor Rst2, which translocates from the nucleus to the cytoplasm upon PKA activation. We found that spPKA-KTR1.0 translocates between the nucleus and cytoplasm in a cAMP-PKA pathway-dependent manner, indicating that the spPKA-KTR1.0 is a reliable indicator of the PKA activity in fission yeast cells. In addition, we implemented a system that simultaneously visualizes and manipulates the cAMP-PKA signaling dynamics by introducing bPAC, a photoactivatable adenylate cyclase, in combination with spPKA-KTR1.0. This system offers an opportunity for investigating the role of the signaling dynamics of the cAMP-PKA pathway in fission yeast cells with higher temporal resolution.

**Take Away:**

- spPKA-KTR1.0 allows visualization of PKA activity at the single-cell level
- Live-cell imaging reveals the transient decrease in PKA activity after M-phase
- Optogenetics allows simultaneous visualization and manipulation of PKA activity

## Introduction

Cells harness nutrient-sensing mechanisms to adapt to the external conditions by controlling their proliferation. In the fission yeast *Schizosaccharomyces pombe* (*S. pombe*), the cyclic 3’-5’ adenosine monophosphate (cAMP)-protein kinase A (PKA) signaling pathway is responsible for sensing glucose, which is one of the preferred carbon sources for *S. pombe* (Hoffman 2005; Hoffman and Winston 1990). Extracellular glucose is primarily detected by a G protein-coupled receptor, Git3, at the plasma membrane, which in turn activates adenylate cyclase, Cyr1, via heterotrimeric G proteins consisting of Gpa2 (Gα), Git5 (Gβ), and Git11 (Gγ) subunits, resulting in the cAMP production (Fig. 1A) (Hoffman and Winston 1991; Maeda, Mochizuki, and Yamamoto 1990; Young et al. 1989; Yamawaki-Kataoka et al. 1989; Isshiki et al. 1992; Welton and Hoffman 2000; Kim et al. 1996; Sheila Landry et al. 2000; S. Landry and Hoffman 2001). The cAMP activates the catalytic subunit of PKA, Pka1, by releasing Pka1 from the inhibitory regulatory subunit, Cgs1 (Gupta et al. 2011a; DeVoti et al. 1991; Maeda et al. 1994). In contrast, a cAMP phosphodiesterase, Cgs2, negatively regulates the PKA activity by degrading cAMP (DeVoti et al. 1991; Mochizuki and Yamamoto 1992). The cAMP-PKA pathway plays a crucial role in divergent cellular processes in fission yeast such as stress resistance (Matsuo and Kawamukai 2017), sexual differentiation (Maeda et al. 1994), spore germination (Hatanaka and Shimoda 2001; Sakai et al. 2023), gluconeogenesis (Hoffman and Winston 1991), cell size control (Kelkar and Martin 2015; Uysal Özdemir et al. 2024), heterochromatin formation (Bao et al. 2022), and autophagy (Pérez-Díaz et al. 2023). Among these, it is well known that cAMP-PKA pathway represses the expression of Ste11, a key transcription factor for sexual differentiation, by inactivating the zinc finger transcription factor Rst2 (Otsubo and Yamamoto 2012). When the nitrogen source is depleted, Ste11 is induced in an Rst2-dependent manner (Higuchi, Watanabe, and Yamamoto 2002), but it is not known how the regulation of PKA activity is coupled with the nitrogen starvation.

**Fig. 1.**
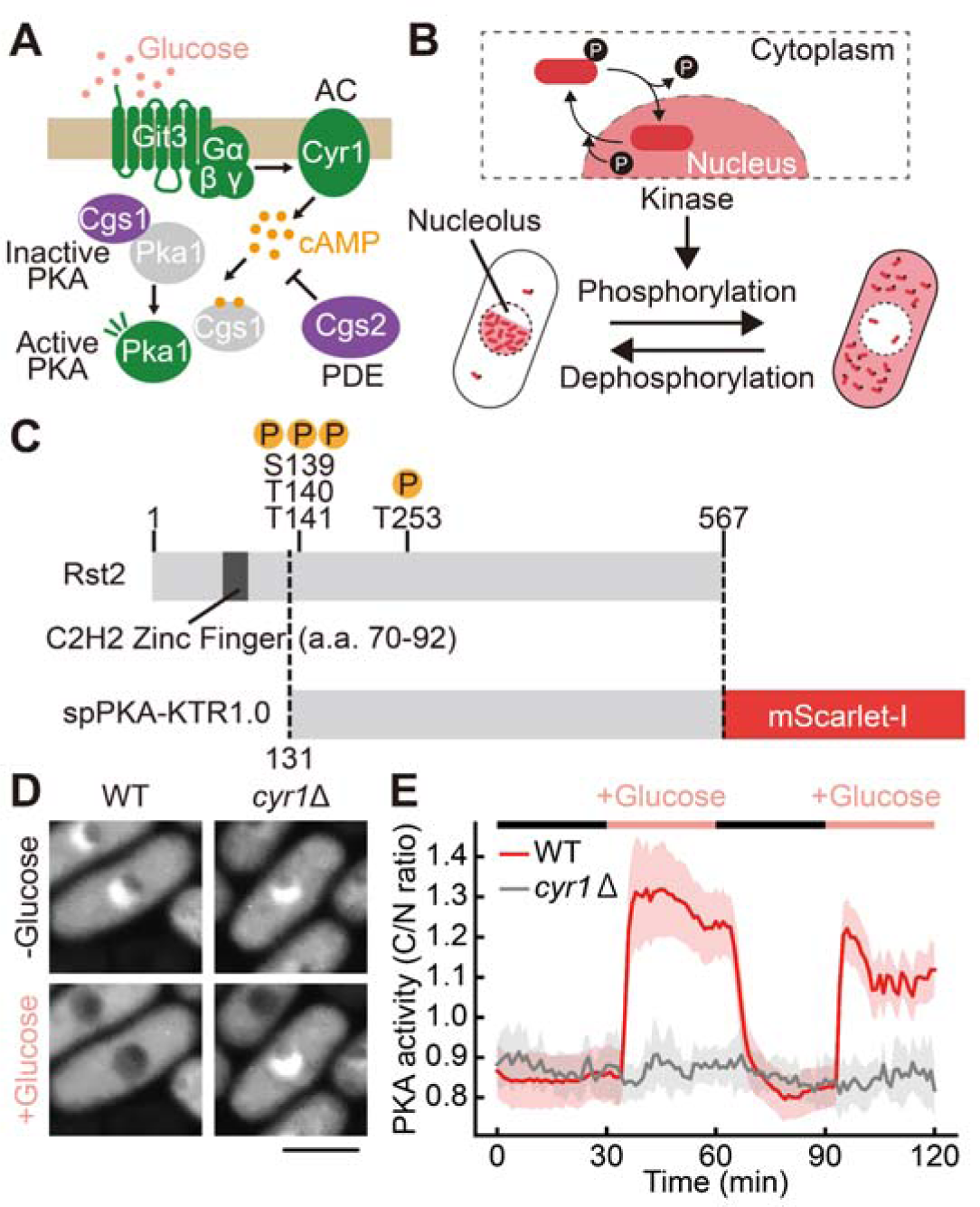
Development of the PKA biosensor spPKA-KTR1.0 for fission yeast cells. (A) Schematic representation of the cAMP-PKA signaling pathway in fission yeast cells. AC, adenylate cyclase; PDE, cAMP phosphodiesterase. (B) Schematic illustration of the phosphorylation-dependent translocation of the kinase translocation reporters (KTRs). Unphosphorylated KTR-based biosensors are primarily localized to the nucleus, whereas upon the kinase activation, the biosensors are phosphorylated and translocated to the cytoplasm. The PKA biosensor, spPKA-KTR1.0, was excluded from the nucleolus. (C) Structure of the KTR-based PKA biosensor spPKA-KTR1.0 (bottom), which consists of the protein region of Rst2 (131–567 a.a., top) and mScarlet-I. To avoid affecting the gene expression, this biosensor does not contain the two Cys_2_His_2_-type zinc finger motifs in the N-terminal region of Rst2 (Kunitomo et al. 2000). The amino acids (S139, T140, T141, T253) in Rst2 are the putative phosphorylation target sites of PKA (Higuchi, Watanabe, and Yamamoto 2002). (D) Representative confocal fluorescence images of wild-type (WT, left) and *cyr1*Δ (right) strains expressing the spPKA-KTR1.0 before (top) and after (bottom) glucose stimulation. Scale bar, 5 μm. (E) Glucose-stimulated PKA activation in wild-type (red, n = 67 cells) and *cyr1*Δ (gray, n = 10 cells) strains expressing the spPKA-KTR1.0. Cells were cultured in 2% glycerol supplemented YEA medium for 16 hours and then stimulated with 2% glucose supplemented YEA medium (+ Glucose). The cytoplasmic-to-nuclear (C/N) ratio of spPKA-KTR1.0 was calculated and the mean values of the C/N ratio are shown with error bars (SD).

The activity of the cAMP-PKA pathway has hitherto been assessed by mating or sporulation efficiency, which is repressed by PKA in fission yeast cells (DeVoti et al. 1991; Mochizuki and Yamamoto 1992; Kawamukai et al. 1991; Maeda, Mochizuki, and Yamamoto 1990; Maeda et al. 1994; Isshiki et al. 1992). Alternatively, PKA activity has been measured by a transcriptional reporter using *fbp1*, a gene encoding fructose-l,6-bisphosphatase, whose transcription is enhanced when PKA is inactivated (Hoffman and Winston 1990, 1991; Nocero et al. 1994; Welton and Hoffman 2000). Biochemical analyses have also been used to investigate intracellular cAMP levels in fission yeast cells (Mochizuki and Yamamoto 1992; DeVoti et al. 1991; Hoffman and Winston 1991; Isshiki et al. 1992; Wang et al. 2005), adenylate cyclase activity *in vitro* (Kawamukai et al. 1991), and PKA activity *in vitro* and *in vivo* by using kemptide (DeVoti et al. 1991) and its endogenous target (Higuchi, Watanabe, and Yamamoto 2002) as substrates. Although these conventional approaches have allowed the analysis of the molecular pathways and genetic interactions involved in the cAMP-PKA pathway, it is still technically challenging to quantitatively monitor the temporal dynamics of PKA activity in a living fission yeast cell.

Genetically encoded fluorescent biosensors are promising tools for the real-time monitoring of signaling dynamics at the single-cell level (Maryu et al. 2018). Förster Resonance Energy Transfer (FRET)-based biosensors are among the most widely used for the visualization of intracellular signaling. The FRET biosensors for cAMP levels and PKA activity were originally developed for mammalian cells (J. Zhang et al. 2001; Komatsu et al. 2011; Ponsioen et al. 2004). Recently, the cAMP biosensors (e.g., yeast-EPAC, camp-EPAC2) (Colombo et al. 2017; Botman et al. 2021) and PKA biosensors (e.g., AKAR3, AKAR3EV, AKAR4) (Colombo et al. 2017, 2022; Botman et al. 2023; Molin et al. 2020) have been successfully adapted for the use in the budding yeast *Saccharomyces cerevisiae*. However, FRET-based biosensors for cAMP and PKA have not yet been applied to the fission yeast *S. pombe*, and therefore it is still unclear when and how PKA activation dynamics commit cell fate decisions. Here, we report tools for the visualization and optical manipulation of PKA activity in a fission yeast cell. We first applied the FRET biosensors for cAMP levels and PKA activity, but these biosensors did not work in *S. pombe* for unknown reasons. Therefore, we developed a biosensor based on the principle of the kinase translocation reporter (KTR) (Regot et al. 2014; Maryu, Matsuda, and Aoki 2016; Miura et al. 2018) to quantitatively monitor the signaling dynamics of the PKA in a living fission yeast cell. By using a blue light-responsive optogenetic tool in combination with the KTR-based PKA biosensor, we succeeded in visualizing and manipulating PKA activity.

## Materials and Methods

### Plasmids

All plasmids used in this study are summarized in Table 1, along with Benchling links including the plasmid sequences and maps. To construct pMNATZA1-spPKA-KTR1.0, the cDNA sequence of Rst2 (131–567 a.a.) was fused to the cDNA sequence of mScarlet-I by PCR and subcloned into pMNATZA1 using Gibson assembly with NEBuilder HiFi DNA Assembly (New England Biolabs, Ipswich, MA). The spPKA-KTR1.0 is stably expressed under the *Padh1* promoter from the Z locus, which is adjacent to the *zfs1* gene on chromosome 2 (Sakuno, Tada, and Watanabe 2009). To construct pMNATZA1-spPKA-KTR1.0-4A, the cDNA sequence of Rst2 (131–567 a.a.) with four mutations (S139A, T140A, T141A, T253A) was amplified, fused to the cDNA sequence of mScarlet-I by PCR and subcloned into pMNATZA1. For the construction of plasmids with truncated regions of Rst2 (pMNATZA1-spPKA-KTR1.1, pMNATZA1-spPKA-KTR1.2, pMNATZA1-spPKA-KTR1.3, and pMNATZA1-spPKA-KTR1.4), the DNA sequences of Rst2 (131–456 a.a., 131–356 a.a., 131–256 a.a., and 131–228 a.a.) were amplified, fused to the cDNA sequence of mScarlet-I by PCR and subcloned into pMNATZA1. To construct pHBCA21-bPAC_sp_opt, the cDNA sequence of bPAC optimized for fission yeast codon usage and the cDNA sequence of the *Padh21* promoter were subcloned into pHBCA1 using conventional ligation with Ligation high Ver.2 (TOYOBO, Osaka, Japan). The amino acid sequence of bPAC was obtained from pGEM-HE-h_bPAC_cmyc (Addgene plasmid #28134) (Stierl et al. 2011). The bPAC is stably expressed under the *Padh21* promoter from the C locus, which is adjacent to the *SPAC26F1.12c* gene on chromosome 1 (Sakuno, Tada, and Watanabe 2009). To construct pMNATZA1-AKAR3EV, the cDNA sequence of AKAR3EV from pCSIIbsr-AKAR3EV (Addgene plasmid #170309) (Komatsu et al. 2011) was subcloned into pMNATZA1 by conventional ligation. To construct the variant pMNATZA1-AKAR3EV-TA, a mutation (T506A) was introduced into pMNATZA1-AKAR3EV. To construct the pMNATZA1-EPACsensor-2051-ApaItoClaI, the cDNA sequence of CFP-Epac-YFP (Nakamoto et al. 2021) was subcloned into pMNATZA1 and the ApaI recognition sequence was replaced by the ClaI recognition sequence.

**Table 1.**
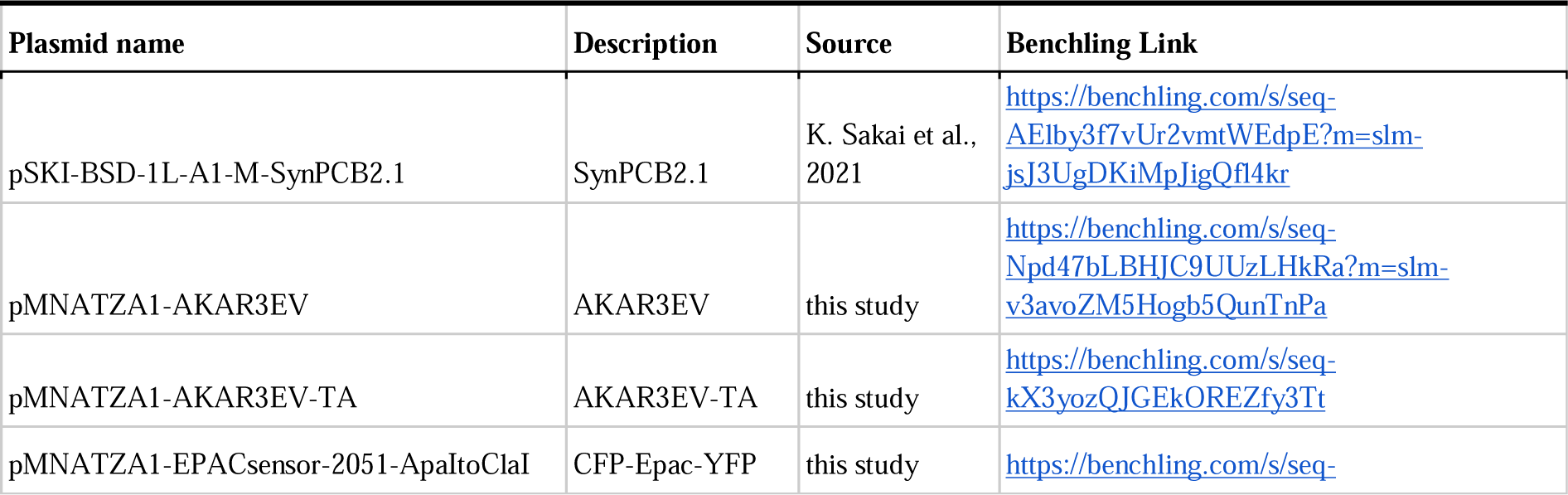
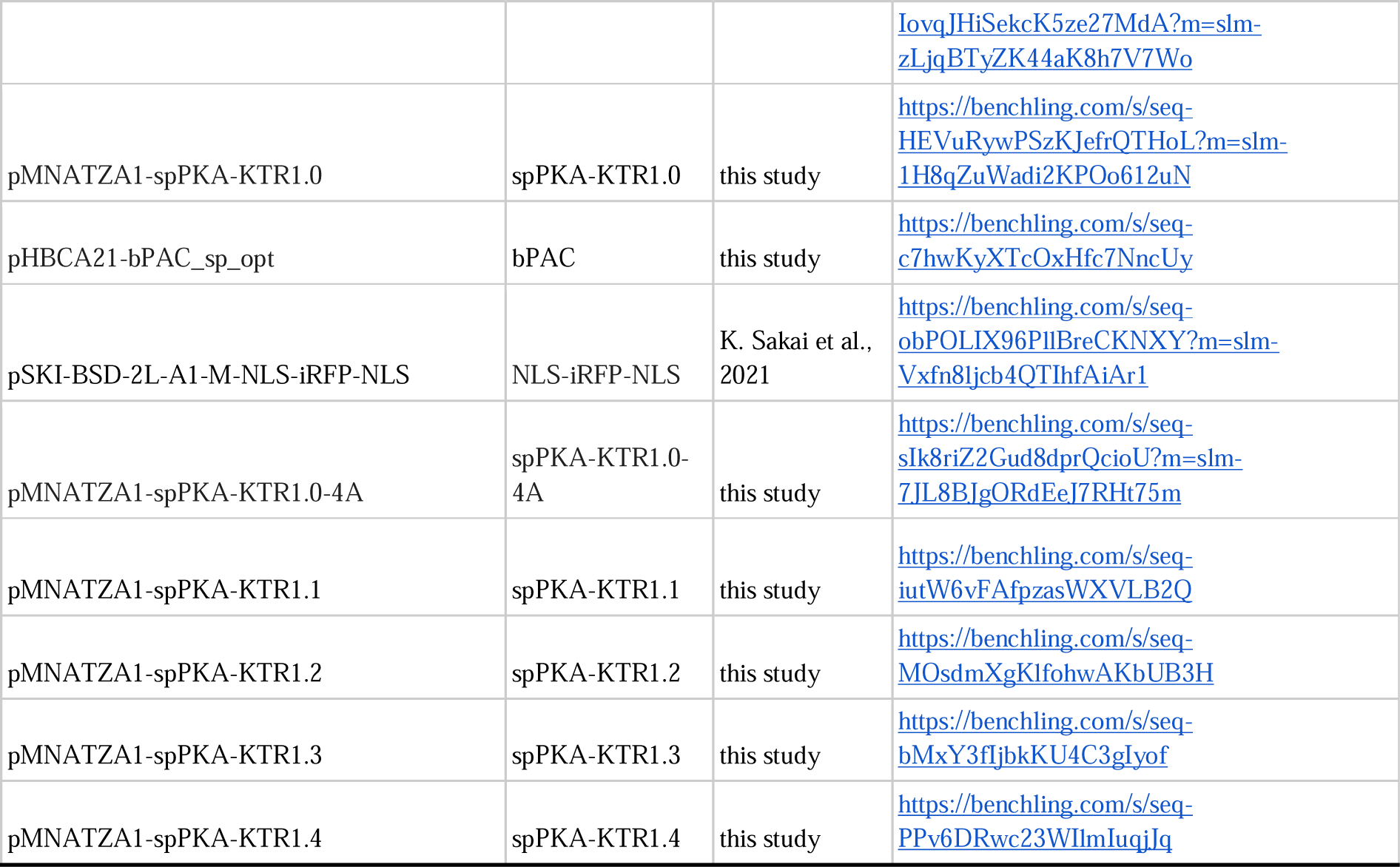
Plasmid list.

### Fission yeast strains and culture conditions

All fission yeast strains used in this study are summarized in Table 2 along with their origins. Unless otherwise noted, the standard methods and media selection followed those previously described (Moreno, Klar, and Nurse 1991). Fission yeast cells were cultured in YEA medium supplemented with high (2%, 111 mM), low (0.2%, 11 mM), or no (0%) glucose. To adjust the osmolyte concentration in the low-glucose medium (glucose concentration of 0 and 0.2%), 2% and 1.8% glycerol were added, respectively. The fission yeast transformation protocol was modified from a previously reported one (Suga and Hatakeyama 2005). To use iRFP in fission yeast, the SynPCB system (Uda et al. 2017, 2020) was further introduced into the cells. Briefly, iRFP requires a chromophore such as biliverdin (BV) and phycocyanobilin (PCB) for emitting fluorescence, but fission yeast *S. Pombe* does not contain genes synthesizing such chromophores. SynPCB is the biosynthesis system consisting of genes from cyanobacteria, efficiently producing PCB and allowing iRFP fluorescence imaging in fission yeast cells (Sakai et al. 2021).

**Table 2.** Schizosaccharomyces pombe strain list.

| Strain name | Genotype | Fig. | Source |
| --- | --- | --- | --- |
| L972 | h- |  | NBRP |
| L975 | h+ |  | NBRP |
| YG038 | h- hta1-mNeonGreen<<kan |  | this study, L972 |
| YG509 | h- 1L::Padh1-SynPCB2.1<<bsd |  | this study, L972 |
| SK014 | h- Z::Padh1-AKAR3EV<<nat | Fig. S1A, S1B | this study, L972 |
| SK023 | h- pka1::kan |  | this study, L972 |
| SK029 | h+ cyr1::kan |  | this study, L975 |
| SK034 | h+ cyr1::kan Z::Padh1-EPACsensor-2051<<nat | Fig. S1C, S1D | this study, SK029 |
| SK045 | h+ cgs2::bsd |  | this study, L975 |
| SK076 | h- cgs1::kan |  | this study, L972 |
| SK077 | h+ cgs1::kan |  | this study, L975 |
| SK094 | h- pka1::kan z::Padh1-AKAR3EV<<nat | Fig. S1A, S1B | this study, SK023 |
| SK095 | h- cgs1::kan z::Padh1-AKAR3EV<<nat | Fig. S1A, S1B | this study, SK076 |
| SK096 | h- z::Padh1-AKAR3EV-TA<<nat | Fig. S1A, S1B | this study, L972 |
| SK097 | h- cgs1::kan z::Padh1-AKAR3EV-TA<<nat | Fig. S1A, S1B | this study, SK076 |
| SK104 | h- pka1::kan z::Padh1-AKAR3EV-TA<<nat | Fig. S1A, S1B | this study, SK023 |
| SK112 | h+ z::Padh1-EPACsensor-2051<<nat | Fig. S1C, S1D | this study, L975 |
| SK113 | h+ cgs2::bsd z::Padh1-EPACsensor-2051<<nat | Fig. S1C, S1D | this study, SK045 |
| SK186 | h- hta1-mNeonGreen<<kan z::Padh1-spPKA-KTR1.0(mScarlet-I)<<nat | Fig. 1D, 1E, Fig. 2A, 2B, 2C, 2D, 2F, Fig. 3A, 3B, Fig. 4A, 4B, 4C, 4D, 4E, Fig. 5A, 5B, 5C, 5D, 5E, Fig. S2A, S2B, Fig. S3, Fig. S4, Movie S1 | this study, YG038 |
| SK188 | h+ cyr1::kan z::Padh1-spPKA-KTR1.0(mScarlet-I)<<nat | Fig. 1D, 1E, Fig. 2A, 2B | this study, SK029 |
| SK192 | h+ cgs2::bsd z::Padh1-spPKA-KTR1.0(mScarlet-I)<<nat | Fig. 2A, 2B | this study, SK045 |
| SK193 | h+ cgs1::kan z::Padh1-spPKA-KTR1.0(mScarlet-I)<<nat | Fig. 2A, 2B, 2C, 2D, 2F, 2G, Fig. 3B | this study, SK077 |
| SK194 | h- pka1::kan z::Padh1-spPKA-KTR1.0(mScarlet-I)<<nat | Fig. 2A, 2B, 2C, 2D, 2F, 2G, Fig. 3B | this study, SK023 |
| SK210 | h- 1L::Padh1-SynPCB2.1<<bsd c::Padh21-bPAC_sp_opt<<hyg |  | this study, YG509 |
| SK211 | h+ cyr1::kan z::Padh1-spPKA-KTR1.0(mScarlet-I)<<nat 2L::Padh1-NLS-iRFP-NLS<<bsd |  | this study, SK188 |
| SK230 | h+ 1L::Padh1-SynPCB2.1<<bsd c::Padh21-bPAC_sp_opt<<hyg cyr1::kan z::Padh1-spPKA-KTR1.0(mScarlet-I)<<nat 2L::Padh1-NLS-iRFP-NLS<<bsd | Fig. 6B, 6C, 6D, Movie S2 | this study, SK210×211 |
| SK261 | h+ cgs1::kan z::Padh1-spPKA-KTR1.0-4A<<nat | Fig. 2C, 2D | this study, SK077 |
| SK264 | h- pka1::kan z::Padh1-spPKA-KTR1.0-4A<<nat | Fig. 2C, 2D | this study, SK023 |
| SK270 | h- hta1-mNeonGreen-kan z::Padh1-spPKA-KTR1.0-4A<<nat | Fig. 2C, 2D, Fig. 3B, Fig. 4B, 4D, 4E, Fig. 5B, 5D, 5E | this study, YG038 |
| SK306 | h- hta1-mNeonGreen<<kan z::Padh1-spPKA-KTR1.1<<nat | Fig. 2F | this study, YG038 |
| SK307 | h- hta1-mNeonGreen<<kan z::Padh1-spPKA-KTR1.2<<nat | Fig. 2F | this study, YG038 |
| SK308 | h- hta1-mNeonGreen<<kan z::Padh1-spPKA-KTR1.3<<nat | Fig. 2F | this study, YG038 |
| SK309 | h- hta1-mNeonGreen<<kan z::Padh1-spPKA-KTR1.4<<nat | Fig. 2F | this study, YG038 |
| SK310 | h+ cgs1::kan z::Padh1-spPKA-KTR1.1<<nat | Fig. 2F, 2G | this study, SK077 |
| SK311 | h+ cgs1::kan z::Padh1-spPKA-KTR1.2<<nat | Fig. 2F, 2G | this study, |
|  |  |  | SK077 |
| SK312 | h+ cgs1::kan z::Padh1-spPKA-KTR1.3<<nat | Fig. 2F, 2G | this study,<br>SK077 |
| SK313 | h+ cgs1::kan z::Padh1-spPKA-KTR1.4<<nat | Fig. 2F, 2G | this study,<br>SK077 |
| SK314 | h- pka1::kan z::Padh1-spPKA-KTR1.1<<nat | Fig. 2F, 2G | this study,<br>SK023 |
| SK315 | h- pka1::kan z::Padh1-spPKA-KTR1.2<<nat | Fig. 2F, 2G | this study,<br>SK023 |
| SK316 | h- pka1::kan z::Padh1-spPKA-KTR1.3<<nat | Fig. 2F, 2G | this study,<br>SK023 |
| SK317 | h- pka1::kan z::Padh1-spPKA-KTR1.4<<nat | Fig. 2F, 2G | this study,<br>SK023 |

### Live-cell fluorescence imaging of fission yeast cells

Fission yeast cells were imaged using an IX83 inverted microscope (Olympus) equipped with an sCMOS camera (ORCA-Fusion BT; Hamamatsu Photonics), a spinning disk confocal unit (CSU-W1; Yokogawa Electric Corporation), an oil immersion objective lenses (UPLXAPO 60X, NA = 1.42, WD = 0.15 mm; Olympus), and diode lasers at wavelengths of 445 nm, 488□nm, 561□nm, and 640□nm. The excitation laser and fluorescence filter settings were as follows: Excitation laser, 445 nm for Turquoise2-GL and mTFP, 488 nm for mNeonGreen, 561□nm for mScarlet-I, and 640□nm for iRFP; excitation dichroic mirrors, DM405/488/561/640 for mNeonGreen, mScarlet-I, and iRFP and DM445/514/640 for Turquoise2-GL, mTFP, YPet, and mVenus; emission filters, 482/35 for Turquoise2-GL and mTFP, 525/50 for mNeonGreen, YPet, and mVenus, 617/73 for mScarlet-I, and 685/40 for iRFP (Yokogawa Electric Corporation). The microscope was controlled by MetaMorph software (ver. 7.10.3). To illuminate cells expressing bPAC with the blue light, blue LED light (450□nm) (LED-41VIS450; OptoCode Corp.) was used from the top of the stage.

For snapshot imaging, fission yeast cells were concentrated by centrifugation at 860 g, mounted on a slide glass (thickness, 0.9–1.2 mm; Matsunami), and sealed with a glass coverslip (thickness, 0.13–0.17 mm; Matsunami). For time-lapse imaging, fission yeast cells were observed by using the CellASIC ONIX2 Microfluidic platform (Merck). The cell suspension was loaded into a trapping chamber (Y04T-04 or Y04C-02) by applying a pressure of 8 psi for 15 seconds. During the time-lapse imaging, the medium was perfused by applying a pressure of 1 psi and the cells were incubated at 32□. During the medium exchange, the new medium was perfused at 8 psi pressure for 15 seconds for washout and then perfused at 1 psi pressure.

For the glucose dose-dependent response of spPKA-KTR1.0 in Figure 3, cells were cultured in YEA medium containing 3% glucose overnight, transferred to low-glucose medium (YEA medium containing 0.1% glucose and 3% glycerol), and then cultured at 32□ for 6 hours. Cells were collected by centrifugation, suspended in YEA medium containing various concentrations of glucose (0.001, 0.003, 0.01, 0.03, 0.1, 0.3, 0.9, 1.5, or 3.0%), and cultured at 32□ for 5 minutes. Cells were then fixed with ice-cold methanol and stored at −30□ overnight. After removal of methanol, cells were resuspended in DDW and imaged as described above.

### Images and data analysis

All imaging data were analyzed and quantified using Fiji/ImageJ (https://fiji.sc/) (Schindelin et al. 2012) and Python. For all images, the background was subtracted using the rolling ball method (rolling ball radius, 50.0 pixels) adopted in Fiji. For the quantification of snapshot images, two regions of interest (ROIs) appropriate for nucleus and cytoplasm were manually selected, the mean fluorescence signal intensities in these ROIs were measured, and the cytoplasm/nucleus (C/N) ratio values for spPKA-KTR1.0 or spPKA-KTR1.0-4A were calculated by dividing the mean intensity in the cytoplasm by that in the nucleus. For quantification of AKAR3EV and CFP-Epac-YFP, ROIs in the cytoplasm were manually selected, the mean fluorescence signal intensities in these ROIs were measured, and the FRET/CFP ratios and CFP/FRET ratios were calculated for AKAR3EV and CFP-Epac-YFP, respectively. For the segmentation, tracking, and quantification of time-lapse images, we used CellTK (Kudo et al. 2018) to quantify the PKA activity from translocation of spPKA-KTR1.0. Briefly, pre-processing, segmentation, tracking, and post-processing were performed using the CellTK functions as we adapted this method in mammalian cells (Tany et al. 2022). In pre-processing, nuclear images, Hta1-mNG or NLS-iRFP-NLS, were processed with a Gaussian blur filter. In the segmentation process, an adaptive thresholding algorithm was used.

Segmented nuclear labels were tracked using the Linear Assignment Problem (LAP) algorithm (Jaqaman et al. 2008) and then processed using a track_neck_cut function in CellTK. In post-processing, short tracking data were excluded, and the cytoplasmic region was defined as an expanding ring around the nuclear segmentation. Quantified feature values (i.e., mean intensity, nuclear area, etc.) of each tracked cell were exported and cleaned in Python. To visualize the heatmaps in Figure 4D and 5E, PKA activities (C/N ratio) of each representative 26 cells were shown from nuclear division to the next. The timing of nuclear division was determined from the acute change in nuclear morphology based on Hta1-mNG images. For the visualization of Figure 2B, 2D, and 2F, each dot represents the C/N ratio of a single cell. In the boxplots, lines represent the median values, boxes represent the interquartile range and whiskers represent the minimum and maximum values, except for outliers, which are defined as 1.5 times the interquartile range. Data visualization and graph generation were performed using Python 3.10 with the modules Numpy 1.21.3, Pandas 1.3.4, Matplotlib 3.4.3, and Seaborn 0.11.2. In Figure 3B, the glucose dose-dependent response of spPKA-KTR1.0 was fitted by the Hill function (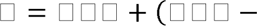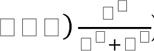), where *n* is the Hill coefficient and *k* is the EC_50_.

**Fig. 2.**
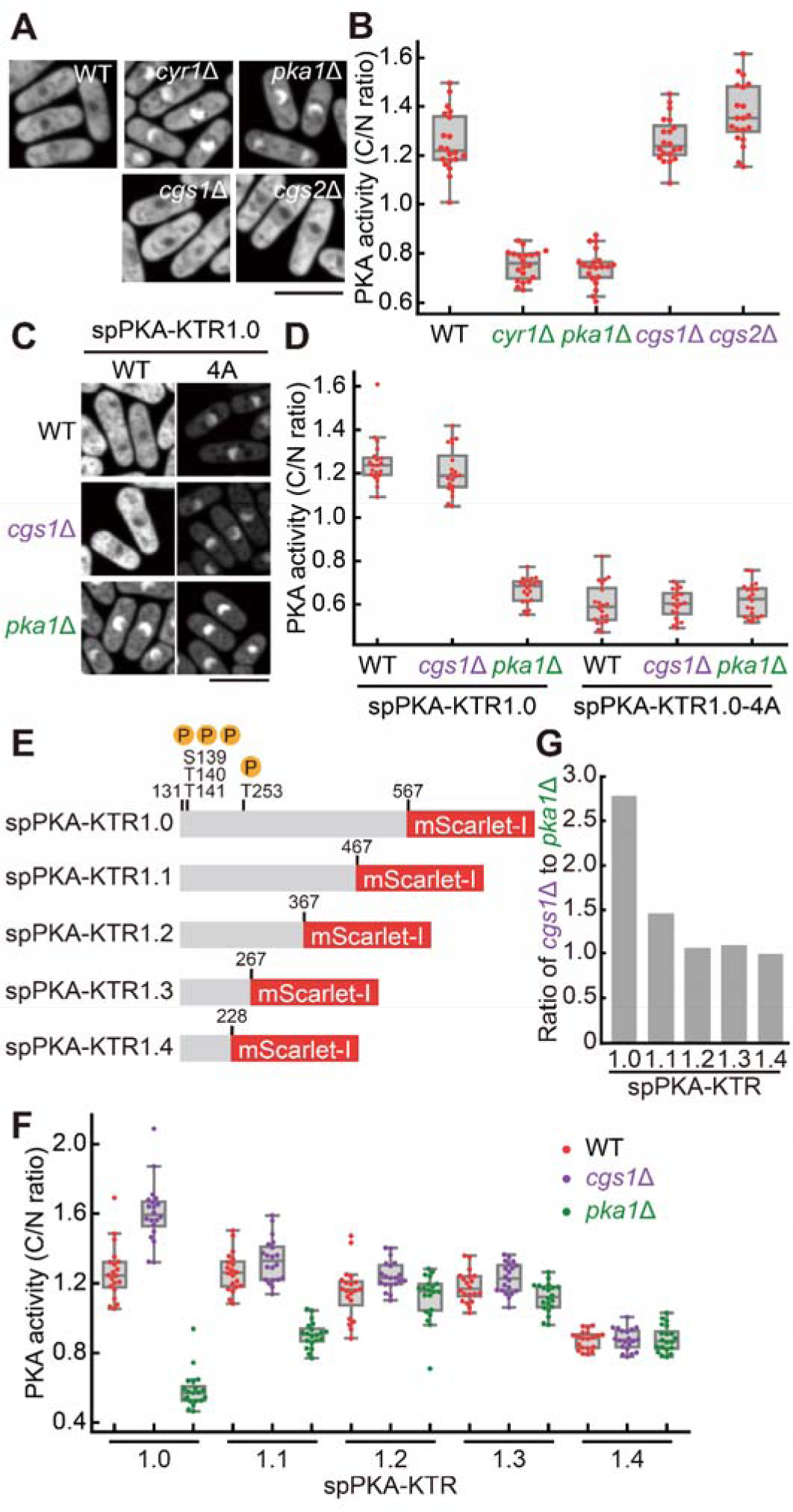
Characterization of the translocation mechanisms of spPKA-KTR1.0. (A) Representative confocal fluorescence images of wild-type (WT), *cyr1*Δ, *pka1*Δ, *cgs1*Δ, and *cgs2*Δ strains expressing the spPKA-KTR1.0 in the 2% glucose-supplemented YEA medium. Scale bar, 10 μm. (B) The PKA activity (C/N ratio) in wild-type (WT), *cyr1*Δ, *pka1*Δ, *cgs1*Δ, and *cgs2*Δ strains expressing the spPKA-KTR1.0 (n = 20 cells). Cells were cultured in YEA medium supplemented with 2% glucose. (C) Representative confocal fluorescence images of wild-type (WT), *cgs1*Δ, and *pka1*Δ strains expressing the spPKA-KTR1.0 or its variant, spPKA-KTR1.0-4A. Scale bar, 10 μm. (D) The PKA activity (C/N ratio) in wild-type (WT), c*gs1*Δ, and *pka1*Δ strains expressing the spPKA-KTR1.0 or its variant, spPKA-KTR1.0-4A (n = 20 cells). Cells were cultured in YEA medium supplemented with 2% glucose. (E) Structure of truncated versions of spPKA-KTR1.0, namely spPKA-KTR1.1–1.4. Each of them has the protein region of Rst2 from 131–467 a.a., 131–367 a.a., 131–267 a.a., and 131–228 a.a., respectively. (F) The C/N ratio of spPKA-KTR1.0–1.4 in wild-type (WT), c*gs1*Δ, and *pka1*Δ strains (n = 20 cells). Cells were grown in YEA medium supplemented with 2% glucose. (G) The ratio of C/N ratio in *cgs1*Δ to that in *pka1*Δ strain expressing the spPKA-KTR1.0–1.4. The ratios were calculated from the data in panel F.

**Fig. 3.**
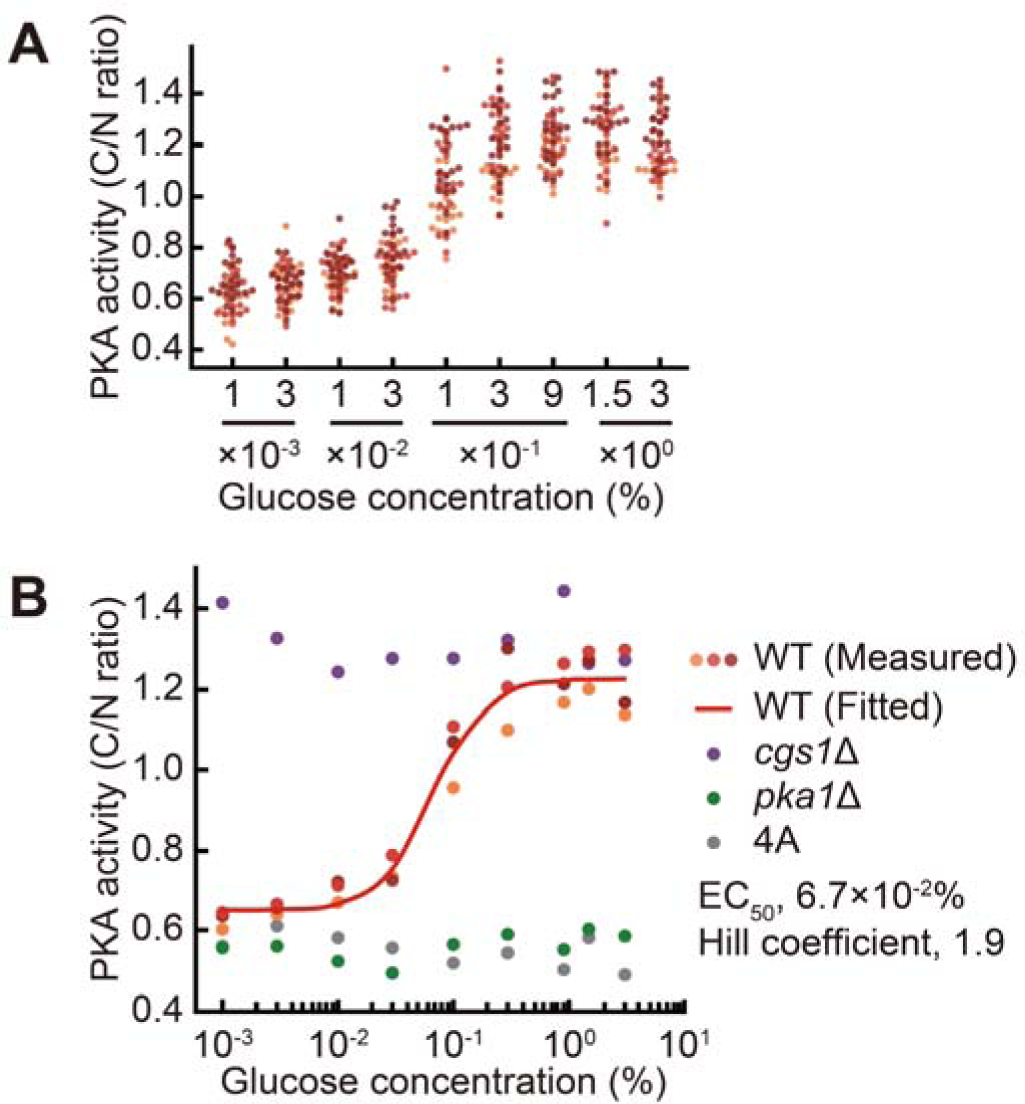
Dose-dependent response of spPKA-KTR1.0 to glucose concentration. (A) PKA activity (C/N ratio) in fission yeast cells expressing spPKA-KTR1.0 in response to various glucose concentrations. Cells were pre-cultured in low-glucose medium (YEA medium supplemented with 0.1% glucose and 3% glycerol), stimulated with YEA medium supplemented with different glucose concentrations (0.001, 0.003, 0.01, 0.03, 0.1, 0.3, 0.9, 1.5, or 3.0%), and then the localization of spPKA-KTR1.0 was observed 5 min after stimulation. The C/N ratios of 20 cells were calculated from each of three independent experiments and plotted with different colored dots. (B) PKA activity (C/N ratio) in wild-type (WT), *cgs1*Δ, and *pka1*Δ strains expressing the spPKA-KTR1.0 or spPKA-KTR1.0-4A (4A) in response to various glucose concentrations (0.001, 0.003, 0.01, 0.03, 0.1, 0.3, 0.9, 1.5, or 3.0%). Each dot represents the median C/N ratio of 20 cells. The solid red line represents the Hill function-based fitted curve for the data from the WT strain expressing spPKA-KTR1.0 (EC_50_, 6.7×10^-2^%; Hill coefficient, 1.9).

**Fig. 4.**
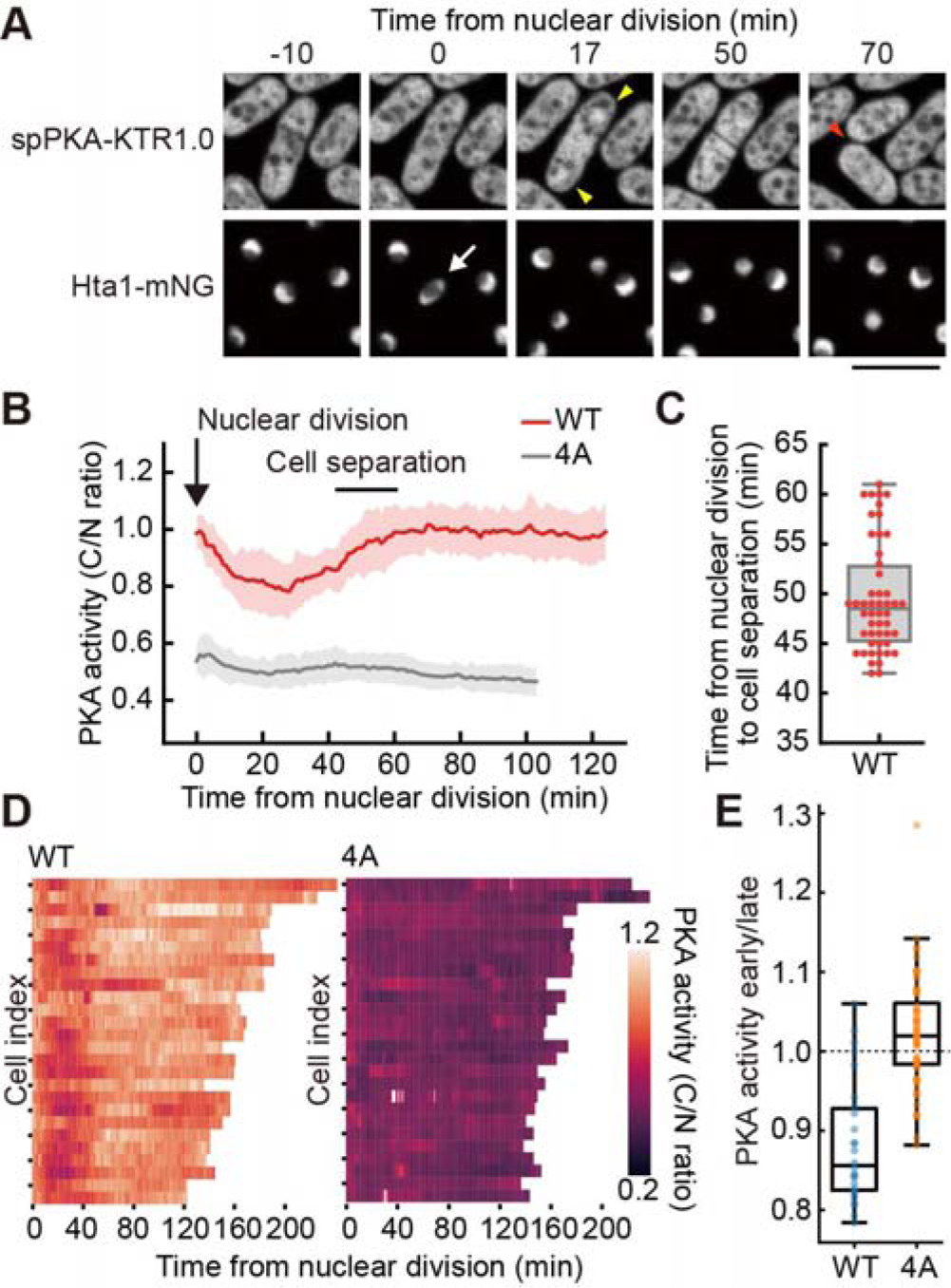
Decreased PKA activity during the cell cycle under high glucose conditions. (A) Representative confocal fluorescence images of the spPKA-KTR1.0 (top) and Hta1 (histone H2A) fused to mNeonGreen (Hta1-mNG, bottom) during nuclear division in wild-type cells. Cells were cultured in YEA medium containing 2% glucose. spPKA-KTR1.0 transiently accumulated in the nucleus during nuclear division in some cells. A white arrow indicates the onset of the nuclear division. Yellow arrows indicate the accumulation of spPKA-KTR1.0 in the nucleus. A red arrow indicates the cell separation. Scale bar, 10 μm. (B) Decreased PKA activity after nuclear division of fission yeast cells cultured in YEA medium containing 2% glucose. The C/N ratios of spPKA-KTR1.0 (WT, n = 26 cells) or spPKA-KTR1.0-4A (4A, n = 64 cells) were calculated, and the mean values of C/N ratios are shown with error bars (SD). Cell separation was indicated by the range of maximum and minimum values quantified in panel C. (C) Quantification of the time from nuclear division to cell separation in cells expressing spPKA-KTR1.0 (n = 50 cells). (D) Heatmap of time courses for PKA activity after nuclear division of fission yeast cells cultured in YEA medium containing 2% glucose. Each row represents a single cell trace of the C/N ratio of spPKA-KTR1.0 (WT, left, n = 26 cells) or spPKA-KTR1.0-4A (4A, right, n = 26 cells). Cells were aligned with the peak of the nuclear area as time 0. (E) To quantify the transient decrease in the C/N ratio in each cell, the averaged C/N ratio during the early time points (0–40 min) was divided by the averaged C/N ratio during later phase (40–80 min). When this value is less than one (dashed line), the C/N ratio exhibits a transient decrease. Most cells (22 cells out of 26 cells) expressing spPKA-KTR1.0 (blue dots) showed the value below one whereas cells expressing spPKA-KTR1.0-4A (orange dots) tended to have values around one (24 cells out of 64 cells were below one).

**Fig. 5.**
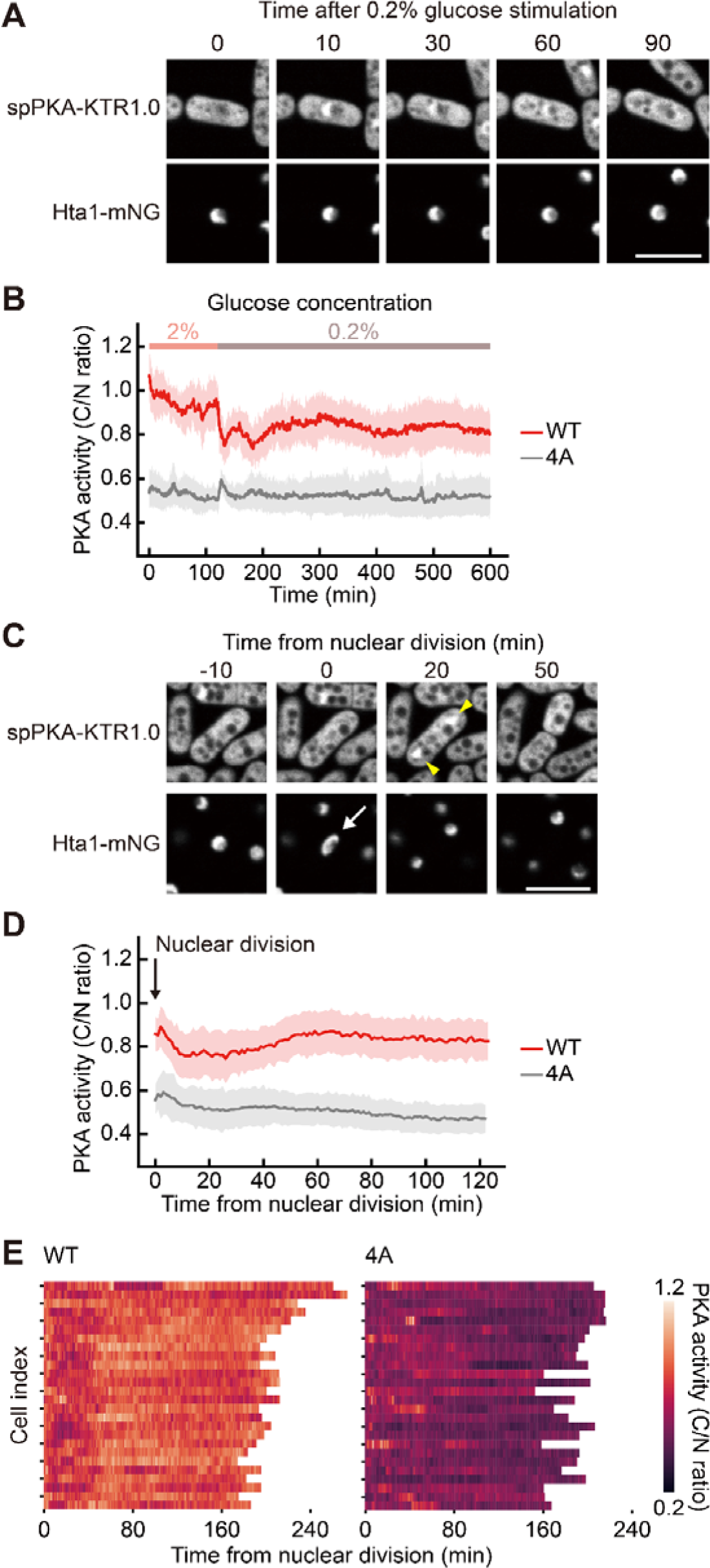
Decreased PKA activity during the cell cycle under glucose-limited conditions. (A) Representative confocal fluorescence images of the spPKA-KTR1.0 (top) and Hta1-mNG (bottom) in wild-type cells after changing the glucose concentration in the medium from 2% to 0.2%. Scale bar, 10 μm. (B) Decreased PKA activity after changing the glucose concentration in the medium from 2% to 0.2% in fission yeast cells. The C/N ratios of spPKA-KTR1.0 (WT, n = 171 cells) or spPKA-KTR1.0-4A (4A, n = 379 cells) were calculated, and the mean values of C/N ratios are shown with error bars (SD). (C) Representative confocal fluorescence images of the spPKA-KTR1.0 (top) and Hta1-mNG (bottom) during nuclear division in wild-type cells. Cells were grown in YEA medium supplemented with 0.2% glucose. A white arrow indicates the onset of the nuclear division. Yellow arrows indicate the accumulation of spPKA-KTR1.0 in the nucleus. Scale bar, 10 μm. (D) Decreased PKA activity after nuclear division of fission yeast cells cultured in YEA medium containing 0.2% glucose. The C/N ratios of spPKA-KTR1.0 (WT, n = 77 cells) or spPKA-KTR1.0-4A (4A, n = 121 cells) were calculated and the mean values of C/N ratios are shown with error bars (SD). (E) Heatmap of the time course of PKA activity after nuclear division of fission yeast cells cultured in YEA medium containing 0.2% glucose. Each row represents a single cell trace of the C/N ratio of spPKA-KTR1.0 (WT, left, n = 26 cells) or spPKA-KTR1.0-4A (4A, right, n = 26 cells). Cells were aligned with the peak of the nuclear area as time 0.

## Results and Discussion

### FRET-based biosensors for PKA activity and cAMP levels did not work in the *S. pombe*

To monitor PKA activity and cAMP levels in the fission yeast *S. pombe*, we first tried the previously reported FRET-based biosensors. Unfortunately, neither the FRET-based the PKA biosensor AKAR3EV (Komatsu et al. 2011) nor the cAMP biosensor CFP-Epac-YFP (Ponsioen et al. 2004) could be used in fission yeast cells (Fig. S1). AKAR3EV is a FRET-based PKA biosensor that was originally developed to measure PKA activity in mammalian cells (Jin Zhang et al. 2005; Komatsu et al. 2011). When AKAR3EV is phosphorylated by PKA, it undergoes conformational changes and exhibits a higher FRET/CFP ratio than in the non-phosphorylated state. As expected, AKAR3EV-TA, in which the threonine residue of PKA phosphorylation is substituted with an alanine residue, showed lower FRET/CFP ratio values than AKAR3EV (Fig. S1A and S1B). However, deletion of either the catalytic subunit *pka1* or the regulatory subunit *cgs1* did not alter the FRET/CFP ratio values of AKAR3EV (Fig. S1A and S1B), suggesting that AKAR3EV is phosphorylated by other kinases.

The FRET-based cAMP biosensor CFP-Epac-YFP (Ponsioen et al. 2004), which is a closed conformation in the absence of cAMP, becomes an open conformation upon binding to cAMP. The increases in cAMP lead to decreases in FRET efficiency. Therefore, we quantified the CFP/FRET ratio of CFP-Epac-YFP as a proxy for cAMP binding. As expected, deletion of the phosphodiesterase gene *cgs2* increased the CFP/FRET values (Fig. S1C and S1D). However, the adenylate cyclase-deficient strain, *cyr1*Δ cells, did not show a significant decrease in the CFP/FRET ratio in fission yeast grown in rich medium (Fig. S1C and S1D). These results suggest that CFP-Epac-YFP is unable to detect the physiological cAMP levels in fission yeast cells due to its lower sensitivity to cAMP than the physiological cAMP levels.

### Development of the KTR-based PKA biosensor, spPKA-KTR1.0, for *S. pombe*

The principle of KTR relies on the concept of converting a phosphorylation event into a nucleocytoplasmic shuttling event. The reporter is exported from the nucleus to the cytoplasm when it is phosphorylated, and imported back into the nucleus when it is dephosphorylated (Fig. 1B). Thus, this type of reporter allows us to estimate kinase activity based on the ratio of cytoplasmic to nuclear fluorescence intensity (C/N ratio). To construct a KTR-based PKA biosensor for fission yeast, we focused on the transcription factor Rst2, which is directly phosphorylated by PKA and exhibits nucleocytoplasmic shuttling in a PKA activity-dependent manner (Fig. 1C) (Higuchi, Watanabe, and Yamamoto 2002; Gupta et al. 2011b; Jiang et al. 2021; Koike et al. 2012). Since overexpression of full-length Rst2 induces growth defects in fission yeast cells (Takenaka et al. 2018; Koike et al. 2012), we deleted the N-terminus of Rst2 up to 130 amino acids (a.a.), which contains the two Cys_2_His_2_-type zinc finger motifs (Kunitomo et al. 2000). The truncated Rst2 fused to mScarlet-I is referred to as spPKA-KTR1.0 (Fig. 1C). We established a fission yeast strain that stably expresses the spPKA-KTR1.0 under the constitutive *adh1* promoter (*Padh1*). The spPKA-KTR1.0 accumulated in the nucleus of fission yeast cells cultured in medium without glucose (Fig. 1D). The reporter rapidly translocated from the nucleus to the cytoplasm when glucose was supplied into the medium (Fig. 1D, Movie S1). The cytoplasmic translocation of the reporter upon glucose stimulation was not observed in adenylate cyclase-deficient cells (*cyr1*Δ) (Fig. 1D). We quantified the C/N ratios of spPKA-KTR1.0, and found that the values started to increase 5 minutes after glucose stimulation and reached a plateau within a minute (Fig. 1E). After glucose washout, the C/N ratios returned to basal levels within 10 minutes (Fig. 1E). In contrast, the C/N ratios of spPKA-KTR1.0 in *cyr1*Δ cells did not change under the same conditions (Fig. 1E). These results support the idea that the nucleocytoplasmic translocation of spPKA-KTR1.0 depends on Cyr1-induced cAMP production and subsequent PKA activation. Compared to the steep decrease in PKA activity with the glucose washout (Fig. 1E), nitrogen starvation did not affect the PKA activity as much (Fig. S2A and B).The spPKA-KTR1.0 was excluded from the nucleolus (Fig. 1B and 1D), suggesting that this reporter may still have some ability to associate with chromatin in the nucleus despite the absence of the zinc finger motifs. A recent study has reported that transcription factors lacking DNA binding domains often localize to chromatin by using their intrinsically disordered regions (IDRs) (Kumar et al. 2023). Rst2 also has IDRs throughout its amino acid sequence, suggesting that spPKA-KTR1.0 preferentially localizes to the chromatin region through IDRs.

### The molecular mechanisms of the spPKA-KTR1.0 translocation in fission yeast cells

We further investigated the mechanisms of the spPKA-KTR1.0 translocation using three approaches. First, we measured the C/N ratio in mutant strains lacking components of the cAMP-PKA pathway (*cyr1*, *pka1*, *cgs1*, or *cgs2*) in the medium containing 2% glucose. The spPKA-KTR1.0 was localized in the cytoplasm in wild-type, *cgs1*Δ, and *cgs2*Δ strains, whereas *cyr1*Δ and *pka1*Δ strains showed nuclear accumulation of spPKA-KTR1.0 (Fig. 2A). The C/N ratios of the *cyr1*Δ and *pka1*Δ strains were reduced compared to that of the wild-type strain, whereas those of the *cgs1*Δ and *cgs2*Δ strains were comparable or higher than that of the wild-type strain (Fig. 2B). These results are consistent with previous findings that Cyr1 and Pka1 are required for PKA activation (Maeda, Mochizuki, and Yamamoto 1990; Maeda et al. 1994) and that PKA is constitutively activated in *cgs1*Δ and *cgs2*Δ strains (DeVoti et al. 1991).

Second, we examined whether PKA-mediated phosphorylation directly controls spPKA-KTR1.0 translocation. Since spPKA-KTR1.0 has two putative PKA target sites in its sequence (Fig. 1C) (Higuchi, Watanabe, and Yamamoto 2002), we substituted the putative clustered phosphorylation sites (S139, T140, T141) and T253 with alanine residues to generate a mutant of spPKA-KTR1.0 (spPKA-KTR1.0-4A). As expected, spPKA-KTR1.0-4A accumulated in the nucleus even in wild-type and *cgs1*Δ strains as well as in the *pka1*Δ strain (Fig. 2C). The cells expressing spPKA-KTR1.0-4A exhibited a lower C/N ratio at the same level as the *pka1*Δ strain expressing spPKA-KTR1.0 (Fig. 2D). These results indicate that the spPKA-KTR1.0 shuttles between the nucleus and cytoplasm in a PKA phosphorylation-dependent manner.

Finally, we examined which regions of spPKA-KTR1.0 are required for its translocation. There seemed to be no nuclear localization signals (NLS) or nuclear export signals (NES) in the amino acid sequence of Rst2. Therefore, for further investigation, we generated a series of truncated spPKA-KTRs, each containing the Rst2 regions 131–467 a.a., 131–367 a.a., 131–267 a.a., and 131– 228 a.a., respectively (Fig. 2E). We quantified the C/N ratio of the truncated spPKA-KTRs in the wild-type, *cgs1*Δ, and *pka1*Δ strains, and found that spPKA-KTR1.1–1.4 did not change their localization in the *cgs1*Δ and *pka1*Δ strains (Fig. 2F). Among the series of spPKA-KTRs, the original spPKA-KTR1.0 showed the highest contrast in C/N ratio between *cgs1*Δ and *pka1*Δ (Fig. 2G), suggesting that multiple domains within the spPKA-KTR1.0 are required for efficient translocation in response to the PKA activity, although we could not find any putative motifs. One possible mechanism for spPKA-KTR1.0 translocation is nuclear export via the 14-3-3 proteins. The 14-3-3 proteins bind to phosphorylated proteins (Muslin et al. 1996) and regulate various cellular functions including nuclear export (Fu, Subramanian, and Masters 2003). Although there are no reports demonstrating a direct interaction between Rst2 and 14-3-3 proteins, growth inhibition in Rst2-overexpressing cells is weakly suppressed by overexpression of 14-3-3 proteins, Rad24 or Rad25, suggesting a genetic interaction between Rst2 and 14-3-3 proteins (Koike et al. 2012). Therefore, nuclear export of phosphorylated Rst2 may be regulated by the Rad24 and Rad25 in a manner similar to the well-studied nuclear export of Cdc25 phosphatase, a cell cycle regulator protein (Lopez-Girona et al. 1999; Kumagai and Dunphy 1999; Dalal et al. 1999).

### Dose-dependent response of spPKA-KTR1.0 to different glucose concentrations in fission yeast cells

To further evaluate whether spPKA-KTR1.0 acts as a reliable biosensor for PKA activity, we measured the dose-dependent response of spPKA-KTR1.0 to a range of glucose concentrations. Cells expressing spPKA-KTR1.0 were cultured in low-glucose medium (YEA medium containing 0.1% glucose and 3% glycerol) and then stimulated with medium containing different glucose concentrations (0.001, 0.003, 0.01, 0.03, 0.1, 0.3, 0.9, 1.5, or 3.0%) (Fig. 3A).The C/N ratio was quantitatively correlated with the glucose concentration, and the EC_50_ and Hill coefficient were 6.7×10^-2^% (3.7 mM) and 1.9, respectively (Fig. 3B). Compared with WT cells, the *cgs1*Δ cells expressing spPKA-KTR1.0 exhibited higher C/N ratios in a glucose concentration-independent manner, whereas the *pka1*Δ cells showed constant low C/N ratios independent of glucose concentration (Fig. 3B). A wide range of C/N ratios within these upper and lower limits were observed in WT cells (Fig. 3B), indicating that the C/N ratios of the *cgs1*Δ cells and *pka1*Δ cells correspond to the upper and lower limits of PKA activity measured by spPKA-KTR1.0. The EC_50_ value (6.7×10^-2^%) is consistent with reports that cells can proliferate normally at glucose concentrations greater than 0.08% (Pluskal et al. 2011; Masuda et al. 2016; Takeda et al. 2015). Moreover, consistent with previous reports on the glucose concentration that determines the transition between proliferation and quiescence (Pluskal et al. 2011), the C/N ratio was close to the basal level upon 0.03% glucose stimulation (Fig. 3B). These results indicate that spPKA-KTR1.0 allows the monitoring of PKA activity under the physiological conditions in fission yeast.

### Visualization of PKA activation dynamics in fission yeast cells

Using the KTR-based PKA biosensor spPKA-KTR1.0, we visualized the dynamics of PKA activity in fission yeast cells under two different conditions. First, the dynamics of PKA activity was measured when cells were cultured in a medium with high glucose concentration (2% glucose), where fission yeast cells grow well. To track the nucleus and cytoplasm over time, endogenous Hta1 (histone H2A) was fused to mNeonGreen (Hta1-mNG) as a nuclear marker. By single-cell tracking of PKA activity (Fig. S3), we found that PKA activity was transiently decreased after nuclear division (Fig. 4A and 4B), and gradually recovered within 40 minutes, coinciding with the cell separation (Fig. 4B and 4C). The transient decrease in PKA activity was observed in most cells (22 cells from 26 cells, Fig. 4D and 4E). Note that the expression level of spPKA-KTR1.0 did not change significantly throughout the cell cycle (Fig. S4). The spPKA-KTR1.0-4A showed a constant low level of the C/N ratio (Fig. 4B and 4D), indicating that the change in the C/N ratio of spPKA-KTR1.0 is regulated by the phosphorylation.

Second, we measured PKA activity in response to acute glucose limitation. In budding yeast, a PKA-regulated transcription factor, Msn2, is known to undergo oscillatory nucleocytoplasmic shuttling in response to the reduction of glucose concentration in the culture medium (Hao et al. 2013). To investigate the dynamics of PKA activity in glucose-limited fission yeast cells, cells expressing spPKA-KTR1.0 were cultured in a high-glucose medium (2% glucose) and then transferred to a low-glucose medium (0.2% glucose). Cell growth was halted soon after the glucose limitation, and spPKA-KTR1.0 was accumulated in the nucleus within 10 min after glucose limitation (Fig. 5A), indicating the reduced PKA activity. Unlike the dynamics of Msn2 in response to glucose in budding yeast cells (Hao et al. 2013), we did not observe the oscillatory nucleocytoplasmic shuttling of spPKA-KTR1.0 in fission yeast cells, suggesting that the dynamics of PKA activity in fission yeast are different from those in budding yeast cells. There was a transient and slight increase in PKA activity immediately after glucose limitation (Fig. 5B). After that, PKA activity gradually recovered over approximately 3 hours, but remained at a lower PKA activity level than before glucose limitation (Fig. 5B). Next, we examined whether PKA activity decreased after nuclear division under the glucose-limited condition (0.2% glucose) as well as the high-glucose condition (2% glucose) (Fig. 4B and 4D). We focused on PKA activity after 280 minutes from 0.2% glucose stimulation (400 min in Fig. 5B), when PKA activity adapted to glucose-limited conditions (Fig. 5B). As in the high-glucose condition, PKA activity was also decreased in the glucose-limited conditions although the transient decrease in the glucose-limited condition was less significant than that in the rich medium due to the low basal PKA level (Fig. 5D and 5E). The similar activation dynamics of PKA reported in mammalian and budding yeast cells using a FRET biosensor and the transcription factor localization, respectively (Vandame et al. 2014; Guerra et al. 2022) suggested a conserved function of PKA in the timely M-phase events.

Although it is known that reduced PKA activity is necessary for sexual differentiation induced by nitrogen depletion (Otsubo and Yamamoto 2012; Higuchi, Watanabe, and Yamamoto 2002), it is not well understood how PKA activity is reduced under these conditions even in the presence of glucose. In this study, we found that during vegetative growth, PKA activity is transiently decreased after nuclear division even in the presence of glucose. If such a transient decrease in PKA activity also occurs during sexual differentiation by arresting cell cycle in G1 phase, the low PKA activity in G1 phase may contribute to the efficient induction of sexual differentiation. The molecular mechanism underlying the reduction of PKA activity from late M to G1 phase is still an open question.

### Manipulation of PKA activity by using photoactivatable adenylate cyclase bPAC

To analyze the causal relationship between intracellular signaling and cellular function, rapid and reversible perturbation of signaling molecules is required. Optogenetics is a promising technique to manipulate signaling molecules and has been widely used to perturb signaling pathways (Goto, Kondo, and Aoki 2021). A photoactivatable adenylate cyclase, bPAC, produces cAMP upon blue light illumination (Iseki et al. 2002; Stierl et al. 2011), which leads to the activation of PKA. Therefore, we implemented bPAC into fission yeast cells to manipulate the signaling dynamics of PKA. We constructed the fission yeast strain expressing bPAC with spPKA-KTR1.0 and NLS-iRFP-NLS (Sakai et al. 2021) to simultaneously manipulate and visualize PKA activity in real time (Fig. 6A). NLS-fused near infrared fluorescent protein, NLS-iRFP-NLS, was used as a nuclear marker for nuclear tracking instead of Hta1-mNG to avoid an unwanted activation of bPAC by the blue light used to excite green fluorescent proteins, such as mNeonGreen. The adenylate cyclase gene, *cyr1*, was deleted to prevent bPAC-independent cAMP production. Under dark conditions, spPKA-KTR1.0 accumulated in the nucleus and was exported from the nucleus to the cytoplasm upon blue light stimulation (Fig. 6B). PKA activity increased within 2 minutes after the blue light illumination, whereas it decreased within 5 minutes after the illumination was turned off (Fig. 6C, Movie S2). Importantly, the level of PKA activity could be fine-tuned by changing the intensity of blue light (Fig. 6D). These results suggest that bPAC-induced PKA activation (∼2 min) is faster than glucose-induced PKA activation (∼5 min) (Fig. 1E). This difference in PKA activation by bPAC and glucose stimulation could be due to three possibilities. One possibility is that glucose stimulation requires time for the glucose signal to be transmitted from the G protein-coupled receptor Git3 to the adenylate cyclase Cyr1, whereas bPAC directly produces cAMP in response to blue light simulation. Second possibility is that the media exchange of glucose stimulation in the microfluidic chamber takes longer than blue light stimulation. The third possibility is that the physiological state is different between glucose-stimulated cells and bPAC-expressing cells because the former have been cultured in the glycerol medium for a long time. Although further investigation is required to determine which possibility is feasible, we found that spPKA-KTR1.0 detects both fast (∼2 min) and slow (∼5 min) activation of PKA. Together, we have confirmed that bPAC can be used to precisely manipulate PKA activity in fission yeast cells with high temporal resolution.

**Fig. 6.**
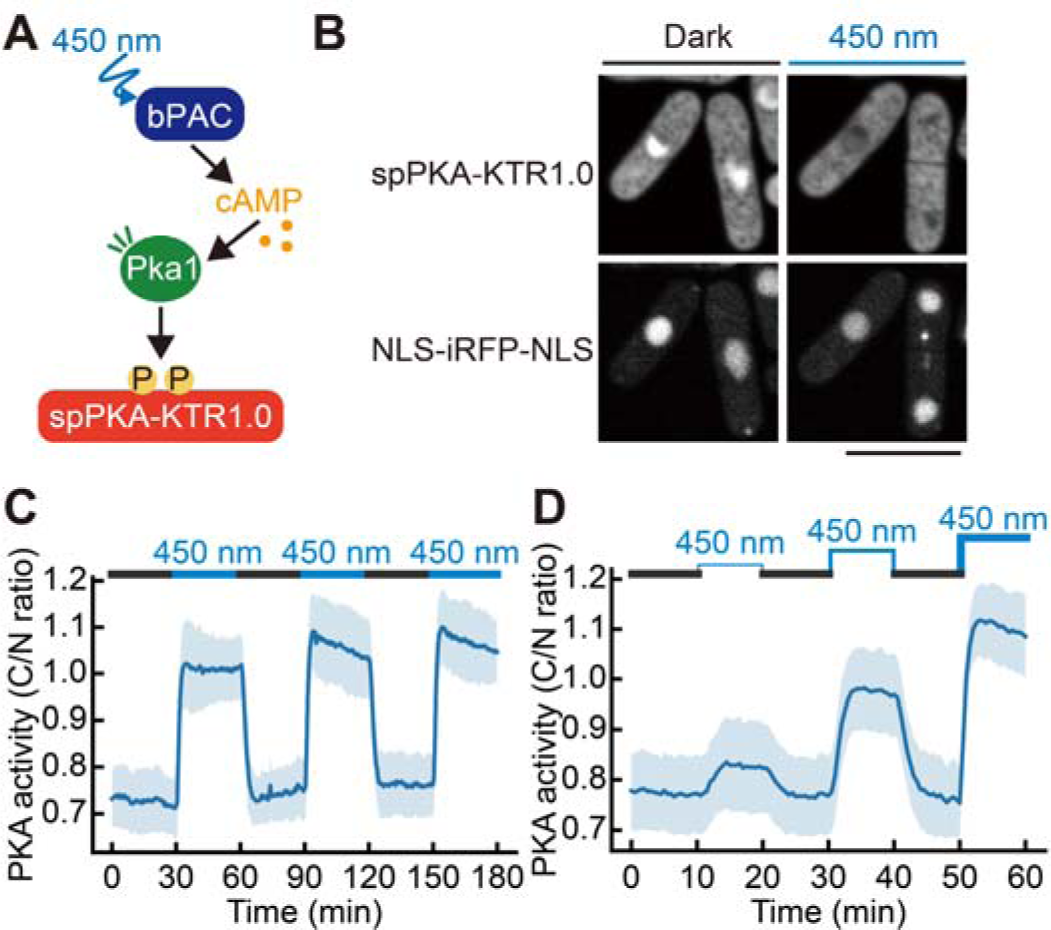
Optogenetic manipulation and visualization of PKA activity by using photoactivatable adenylate cyclase bPAC and spPKA-KTR1.0. (A) Schematic illustration of visualization and manipulation of PKA activity by using spPKA-KTR1.0 and photoactivatable adenylate cyclase bPAC. Upon illumination with blue light (450 nm), bPAC catalyzes the production of cAMP from ATP, resulting in the activation of PKA, which is monitored by spPKA-KTR1.0. (B) Representative confocal fluorescence images of the *cyr1*Δ strain expressing the spPKA-KTR1.0 (top), NLS-iRFP-NLS (bottom), and bPAC. Cells were cultured under dark conditions (left) in 2% glucose-supplemented YEA medium and then stimulated with blue light (450 nm) for 8 min (right). Scale bar, 10 μm. (C and D) Optogenetic control of PKA signaling dynamics in the *cyr1*Δ strain expressing the spPKA-KTR1.0 and bPAC. Due to the deletion of the *cyr1* gene, cAMP production in this strain is dependent on the bPAC. The intensity of blue light illumination was set to constant (C) or stepwise (D). The C/N ratios of spPKA-KTR1.0 were calculated, and the mean values of the C/N ratios are shown with error bars (SD).

## Conclusions

In conclusion, we have developed and characterized the KTR-based PKA biosensor, spPKA-KTR1.0, which is a reliable indicator for the PKA activity in fission yeast cells. Taking advantage of KTR, we have also implemented the photoactivatable adenylate cyclase, bPAC, in fission yeast to simultaneously visualize and manipulate PKA activity with a high temporal resolution. The spPKA-KTR1.0 and bPAC would allow us to investigate the dynamics of PKA activity under different environmental conditions. For example, PKA activity is still required to regulate cell size via microtubule destabilization even under the glucose-limited conditions, where PKA activity is thought to be low (Kelkar and Martin 2015). In addition, PKA activity would be dynamically altered during sexual differentiation, and would be coupled with TORC1 signaling pathway, which mainly senses the nitrogen starvation (Otsubo and Yamamoto 2012). In future studies, measuring and manipulating the dynamics of PKA activity under such conditions will reveal the physiological relevance of the dynamics of the cAMP-PKA pathway throughout the fission yeast life cycle.

## Acknowledgments

We thank all members of the Aoki Laboratory for their helpful discussions and assistance. We also thank members of the Tokai Tor Conference (ToToCo) for the general discussion. Some fission yeast strains were provided by the National Bio-Resource Project (NBRP), Japan. K.A. was supported by JSPS KAKENHI grants (nos. 18H02444, 19H05798, and 22H02625). Y.G. was supported by a JST, ACT-X grant (no. JPMJAX22B8), by JSPS KAKENHI grants (nos.19K16050 and 22K15110), by a Jigami Yoshifumi Memorial Research grant, and by a Sumitomo Research grant. K.S. was supported by a JSPS KAKENHI grant (no. 22J10844) and The Graduate University for Advanced Studies, SOKENDAI (SOKENDAI Student Dispatch Program).

## Author Contribution Statement

**Keiichiro Sakai**: Methodology, Software, Validation, Formal analysis, Investigation, Data Curation, Writing - Original Draft, Writing - Review & Editing, Visualization, Funding acquisition **Kazuhiro Aoki**: Conceptualization, Resources, Writing - Original Draft, Writing - Review & Editing, Supervision, Project administration, Funding acquisition **Yuhei Goto**: Conceptualization, Methodology, Software, Formal analysis, Investigation, Resources, Data Curation, Writing - Original Draft, Writing - Review & Editing, Visualization, Supervision, Project administration, Funding acquisition

## Data Availability Statement

The data that support the findings of this study are openly available in SSDB at http://doi.org/10.24631/ssbd.repos.2024.02.341, reference number ssbd-repos-000341.

## References

Bao, Kehan, Chun-Min Shan, Xiao Chen, Gulzhan Raiymbek, Jeremy G. Monroe, Yimeng Fang, Takenori Toda, et al. 2022. “The cAMP Signaling Pathway Regulates Epe1 Protein Levels and Heterochromatin Assembly.” PLoS Genetics 18 (2): e1010049.

Botman, Dennis, Sineka Kanagasabapathi, Philipp Savakis, and Bas Teusink. 2023. “Using the AKAR3-EV Biosensor to Assess Sch9p- and PKA-Signalling in Budding Yeast.” FEMS Yeast Research 23 (January). 10.1093/femsyr/foad029.

Botman, Dennis, Tom G. O’Toole, Joachim Goedhart, Frank J. Bruggeman, Johan H. van Heerden, and Bas Teusink. 2021. “A Yeast FRET Biosensor Enlightens cAMP Signaling.” Molecular Biology of the Cell 32 (13): 1229–40.

Colombo, Sonia, Serena Broggi, Maddalena Collini, Laura D’Alfonso, Giuseppe Chirico, and Enzo Martegani. 2017. “Detection of cAMP and of PKA Activity in Saccharomyces Cerevisiae Single Cells Using Fluorescence Resonance Energy Transfer (FRET) Probes.” Biochemical and Biophysical Research Communications 487 (3): 594–99.

Colombo, Sonia, Elisa Longoni, Marco Gnugnoli, Stefano Busti, and Enzo Martegani. 2022. “Fast Detection of PKA Activity in Saccharomyces Cerevisiae Cell Population Using AKAR Fluorescence Resonance Energy Transfer Probes.” Cellular Signalling 92 (April): 110262.

Dalal, S. N., C. M. Schweitzer, J. Gan, and J. A. DeCaprio. 1999. “Cytoplasmic Localization of Human cdc25C during Interphase Requires an Intact 14-3-3 Binding Site.” Molecular and Cellular Biology 19 (6): 4465–79.

DeVoti, J., G. Seydoux, D. Beach, and M. McLeod. 1991. “Interaction between ran1+ Protein Kinase and cAMP Dependent Protein Kinase as Negative Regulators of Fission Yeast Meiosis.” The EMBO Journal 10 (12): 3759–68.

Fu, Haian, Romesh R. Subramanian, and Shane C. Masters. 2003. “14-3-3 Proteins: Structure, Function, and Regulation,” November. 10.1146/annurev.pharmtox.40.1.617.

Goto, Yuhei, Yohei Kondo, and Kazuhiro Aoki. 2021. “Visualization and Manipulation of Intracellular Signaling.” Advances in Experimental Medicine and Biology 1293: 225–34.

Guerra, Paolo, Luc-Alban P. E. Vuillemenot, Yulan B. van Oppen, Marije Been, and Andreas Milias-Argeitis. 2022. “TORC1 and PKA Activity towards Ribosome Biogenesis Oscillates in Synchrony with the Budding Yeast Cell Cycle.” Journal of Cell Science 135 (18). 10.1242/jcs.260378.

Gupta, Dipali Rani, Swapan Kumar Paul, Yasuo Oowatari, Yasuhiro Matsuo, and Makoto Kawamukai. 2011a. “Complex Formation, Phosphorylation, and Localization of Protein Kinase A of Schizosaccharomyces Pombe upon Glucose Starvation.” Bioscience, Biotechnology, and Biochemistry 75 (8): 1456–65.

Gupta, Dipali Rani, Swapan Kumar Paul, Yasuo Oowatari, Yasuhiro Matsuo, and Makoto Kawamukai. 2011b. “Multistep Regulation of Protein Kinase A in Its Localization, Phosphorylation and Binding with a Regulatory Subunit in Fission Yeast.” Current Genetics 57 (5): 353–65.

Hao, Nan, Bogdan A. Budnik, Jeremy Gunawardena, and Erin K. O’Shea. 2013. “Tunable Signal Processing through Modular Control of Transcription Factor Translocation.” Science 339 (6118): 460–64.

Hatanaka, M., and C. Shimoda. 2001. “The Cyclic AMP/PKA Signal Pathway Is Required for Initiation of Spore Germination in Schizosaccharomyces Pombe.” Yeast 18 (3): 207–17.

Higuchi, Toru, Yoshinori Watanabe, and Masayuki Yamamoto. 2002. “Protein Kinase A Regulates Sexual Development and Gluconeogenesis through Phosphorylation of the Zn Finger Transcriptional Activator Rst2p in Fission Yeast.” Molecular and Cellular Biology 22 (1): 1– 11.

Hoffman, C. S. 2005. “Glucose Sensing via the Protein Kinase A Pathway in Schizosaccharomyces Pombe.” Biochemical Society Transactions 33 (Pt 1): 257–60.

Hoffman, C. S., and F. Winston. 1990. “Isolation and Characterization of Mutants Constitutive for Expression of the fbp1 Gene of Schizosaccharomyces Pombe.” Genetics 124 (4): 807–16.

Hoffman, C. S., and F. Winston. 1991. “Glucose Repression of Transcription of the Schizosaccharomyces Pombe fbp1 Gene Occurs by a cAMP Signaling Pathway.” Genes & Development 5 (4): 561–71.

Iseki, Mineo, Shigeru Matsunaga, Akio Murakami, Kaoru Ohno, Kiyoshi Shiga, Kazuichi Yoshida, Michizo Sugai, Tetsuo Takahashi, Terumitsu Hori, and Masakatsu Watanabe. 2002. “A Blue-Light-Activated Adenylyl Cyclase Mediates Photoavoidance in Euglena Gracilis.” Nature 415 (6875): 1047–51.

Isshiki, T., N. Mochizuki, T. Maeda, and M. Yamamoto. 1992. “Characterization of a Fission Yeast Gene, gpa2, That Encodes a G Alpha Subunit Involved in the Monitoring of Nutrition.” Genes & Development 6 (12b): 2455–62.

Jaqaman, Khuloud, Dinah Loerke, Marcel Mettlen, Hirotaka Kuwata, Sergio Grinstein, Sandra L. Schmid, and Gaudenz Danuser. 2008. “Robust Single-Particle Tracking in Live-Cell Time-Lapse Sequences.” Nature Methods 5 (8): 695–702.

Jiang, Guanglie, Qiannan Liu, Toshiaki Kato, Hao Miao, Xiang Gao, Kun Liu, Si Chen, Norihiro Sakamoto, Takayoshi Kuno, and Yue Fang. 2021. “Role of Mitochondrial Complex III/IV in the Activation of Transcription Factor Rst2 in Schizosaccharomyces Pombe.” Molecular Microbiology 115 (6): 1323–38.

Kawamukai, M., K. Ferguson, M. Wigler, and D. Young. 1991. “Genetic and Biochemical Analysis of the Adenylyl Cyclase of Schizosaccharomyces Pombe.” Cell Regulation 2 (2): 155–64.

Kelkar, Manasi, and Sophie G. Martin. 2015. “PKA Antagonizes CLASP-Dependent Microtubule Stabilization to Re-Localize Pom1 and Buffer Cell Size upon Glucose Limitation.” Nature Communications 6 (October): 8445.

Kim, D. U., S. K. Park, K. S. Chung, M. U. Choi, and H. S. Yoo. 1996. “The G Protein Beta Subunit Gpb1 of Schizosaccharomyces Pombe Is a Negative Regulator of Sexual Development.” Molecular & General Genetics: MGG 252 (1-2): 20–32.

Koike, Atsushi, Toshiaki Kato, Reiko Sugiura, Yan Ma, Yuki Tabata, Koji Ohmoto, Susie O. Sio, and Takayoshi Kuno. 2012. “Genetic Screening for Regulators of Prz1, a Transcriptional Factor Acting Downstream of Calcineurin in Fission Yeast.” The Journal of Biological Chemistry 287 (23): 19294–303.

Komatsu, Naoki, Kazuhiro Aoki, Masashi Yamada, Hiroko Yukinaga, Yoshihisa Fujita, Yuji Kamioka, and Michiyuki Matsuda. 2011. “Development of an Optimized Backbone of FRET Biosensors for Kinases and GTPases.” Molecular Biology of the Cell 22 (23): 4647–56.

Kudo, Takamasa, Stevan Jeknić, Derek N. Macklin, Sajia Akhter, Jacob J. Hughey, Sergi Regot, and Markus W. Covert. 2018. “Live-Cell Measurements of Kinase Activity in Single Cells Using Translocation Reporters.” Nature Protocols 13 (1): 155–69.

Kumagai, A., and W. G. Dunphy. 1999. “Binding of 14-3-3 Proteins and Nuclear Export Control the Intracellular Localization of the Mitotic Inducer Cdc25.” Genes & Development 13 (9): 1067–72.

Kumar, Divya Krishna, Felix Jonas, Tamar Jana, Sagie Brodsky, Miri Carmi, and Naama Barkai. 2023. “Complementary Strategies for Directing in Vivo Transcription Factor Binding through DNA Binding Domains and Intrinsically Disordered Regions.” Molecular Cell 83 (9): 1462– 73.e5.

Kunitomo, H., T. Higuchi, Y. Iino, and M. Yamamoto. 2000. “A Zinc-Finger Protein, Rst2p, Regulates Transcription of the Fission Yeast ste11(+) Gene, Which Encodes a Pivotal Transcription Factor for Sexual Development.” Molecular Biology of the Cell 11 (9): 3205–17.

Landry, Sheila, Maria T. Pettit, Ethel Apolinario, and Charles S. Hoffman. 2000. “The Fission Yeast git5 Gene Encodes a Gβ Subunit Required for Glucose-Triggered Adenylate Cyclase Activation.” Genetics 154 (4): 1463–71.

Landry, S., and C. S. Hoffman. 2001. “The git5 Gbeta and git11 Ggamma Form an Atypical Gbetagamma Dimer Acting in the Fission Yeast glucose/cAMP Pathway.” Genetics 157 (3): 1159–68.

Lopez-Girona, A., B. Furnari, O. Mondesert, and P. Russell. 1999. “Nuclear Localization of Cdc25 Is Regulated by DNA Damage and a 14-3-3 Protein.” Nature 397 (6715): 172–75.

Maeda, T., N. Mochizuki, and M. Yamamoto. 1990. “Adenylyl Cyclase Is Dispensable for Vegetative Cell Growth in the Fission Yeast Schizosaccharomyces Pombe.” Proceedings of the National Academy of Sciences of the United States of America 87 (20): 7814–18.

Maeda, T., Y. Watanabe, H. Kunitomo, and M. Yamamoto. 1994. “Cloning of the pka1 Gene Encoding the Catalytic Subunit of the cAMP-Dependent Protein Kinase in Schizosaccharomyces Pombe.” The Journal of Biological Chemistry 269 (13): 9632–37.

Maryu, Gembu, Michiyuki Matsuda, and Kazuhiro Aoki. 2016. “Multiplexed Fluorescence Imaging of ERK and Akt Activities and Cell-Cycle Progression.” Cell Structure and Function 41 (2): 81–92.

Maryu, Gembu, Haruko Miura, Youichi Uda, Akira T. Komatsubara, Michiyuki Matsuda, and Kazuhiro Aoki. 2018. “Live-Cell Imaging with Genetically Encoded Protein Kinase Activity Reporters.” Cell Structure and Function 43 (1): 61–74.

Masuda, Fumie, Mahiro Ishii, Ayaka Mori, Lisa Uehara, Mitsuhiro Yanagida, Kojiro Takeda, and Shigeaki Saitoh. 2016. “Glucose Restriction Induces Transient G2 Cell Cycle Arrest Extending Cellular Chronological Lifespan.” Scientific Reports 6 (January): 19629.

Matsuo, Yasuhiro, and Makoto Kawamukai. 2017. “cAMP-Dependent Protein Kinase Involves Calcium Tolerance through the Regulation of Prz1 in Schizosaccharomyces Pombe.” Bioscience, Biotechnology, and Biochemistry 81 (2): 231–41.

Miura, Haruko, Yohei Kondo, Michiyuki Matsuda, and Kazuhiro Aoki. 2018. “Cell-to-Cell Heterogeneity in p38-Mediated Cross-Inhibition of JNK Causes Stochastic Cell Death.” Cell Reports 24 (10): 2658–68.

Mochizuki, N., and M. Yamamoto. 1992. “Reduction in the Intracellular cAMP Level Triggers Initiation of Sexual Development in Fission Yeast.” Molecular & General Genetics: MGG 233 (1-2): 17–24.

Molin, Mikael, Katarina Logg, Kristofer Bodvard, Ken Peeters, Annabelle Forsmark, Friederike Roger, Anna Jörhov, et al. 2020. “Protein Kinase A Controls Yeast Growth in Visible Light.” BMC Biology 18 (1): 168.

Moreno, Sergio, Amar Klar, and Paul Nurse. 1991. “[56] Molecular Genetic Analysis of Fission Yeast Schizosaccharomyces Pombe.” In Methods in Enzymology, 194:795–823. Academic Press.

Muslin, A. J., J. W. Tanner, P. M. Allen, and A. S. Shaw. 1996. “Interaction of 14-3-3 with Signaling Proteins Is Mediated by the Recognition of Phosphoserine.” Cell 84 (6): 889–97.

Nakamoto, Chihiro, Yuhei Goto, Yoko Tomizawa, Yuko Fukata, Masaki Fukata, Kasper Harpsøe, David E. Gloriam, Kazuhiro Aoki, and Tomonori Takeuchi. 2021. “A Novel Red Fluorescence Dopamine Biosensor Selectively Detects Dopamine in the Presence of Norepinephrine in Vitro.” Molecular Brain 14 (1): 173.

Nocero, M., T. Isshiki, M. Yamamoto, and C. S. Hoffman. 1994. “Glucose Repression of fbp1 Transcription of Schizosaccharomyces Pombe Is Partially Regulated by Adenylate Cyclase Activation by a G Protein Alpha Subunit Encoded by gpa2 (git8).” Genetics 138 (1): 39–45.

Otsubo, Yoko, and Masayuki Yamamoto. 2012. “Signaling Pathways for Fission Yeast Sexual Differentiation at a Glance.” Journal of Cell Science 125 (Pt 12): 2789–93.

Pérez-Díaz, Armando Jesús, Beatriz Vázquez-Marín, Jero Vicente-Soler, Francisco Prieto-Ruiz, Teresa Soto, Alejandro Franco, José Cansado, and Marisa Madrid. 2023. “cAMP-Protein Kinase A and Stress-Activated MAP Kinase Signaling Mediate Transcriptional Control of Autophagy in Fission Yeast during Glucose Limitation or Starvation.” Autophagy 19 (4): 1311–31.

Pluskal, Tomáš, Takeshi Hayashi, Shigeaki Saitoh, Asuka Fujisawa, and Mitsuhiro Yanagida. 2011. “Specific Biomarkers for Stochastic Division Patterns and Starvation-Induced Quiescence under Limited Glucose Levels in Fission Yeast: Fission Yeast Division under Glucose Starvation.” The FEBS Journal 278 (8): 1299–1315.

Ponsioen, Bas, Jun Zhao, Jurgen Riedl, Fried Zwartkruis, Gerard van der Krogt, Manuela Zaccolo, Wouter H. Moolenaar, Johannes L. Bos, and Kees Jalink. 2004. “Detecting cAMP-Induced Epac Activation by Fluorescence Resonance Energy Transfer: Epac as a Novel cAMP Indicator.” EMBO Reports 5 (12): 1176–80.

Regot, Sergi, Jacob J. Hughey, Bryce T. Bajar, Silvia Carrasco, and Markus W. Covert. 2014. “High-Sensitivity Measurements of Multiple Kinase Activities in Live Single Cells.” Cell 157 (7): 1724–34.

Sakai, Keiichiro, Yohei Kondo, Hiroyoshi Fujioka, Mako Kamiya, Kazuhiro Aoki, and Yuhei Goto. 2021. “Near-Infrared Imaging in Fission Yeast Using a Genetically Encoded Phycocyanobilin Biosynthesis System.” Journal of Cell Science 134 (24). 10.1242/jcs.259315.

Sakai, Keiichiro, Yohei Kondo, Yuhei Goto, and Kazuhiro Aoki. 2023. “Cytoplasmic Fluidization Triggers Breaking Spore Dormancy in Fission Yeast.” bioRxiv. 10.1101/2023.09.27.559686.

Sakuno, Takeshi, Kenji Tada, and Yoshinori Watanabe. 2009. “Kinetochore Geometry Defined by Cohesion within the Centromere.” Nature 458 (7240): 852–58.

Schindelin, Johannes, Ignacio Arganda-Carreras, Erwin Frise, Verena Kaynig, Mark Longair, Tobias Pietzsch, Stephan Preibisch, et al. 2012. “Fiji: An Open-Source Platform for Biological-Image Analysis.” Nature Methods 9 (7): 676–82.

Stierl, Manuela, Patrick Stumpf, Daniel Udwari, Ronnie Gueta, Rolf Hagedorn, Aba Losi, Wolfgang Gärtner, et al. 2011. “Light Modulation of Cellular cAMP by a Small Bacterial Photoactivated Adenylyl Cyclase, bPAC, of the Soil Bacterium Beggiatoa.” The Journal of Biological Chemistry 286 (2): 1181–88.

Suga, Minoru, and Toyomasa Hatakeyama. 2005. “A Rapid and Simple Procedure for High-Efficiency Lithium Acetate Transformation of Cryopreserved Schizosaccharomyces Pombe Cells.” Yeast 22 (10): 799–804.

Takeda, Kojiro, Caroline Starzynski, Ayaka Mori, and Mitsuhiro Yanagida. 2015. “The Critical Glucose Concentration for Respiration-Independent Proliferation of Fission Yeast, Schizosaccharomyces Pombe.” Mitochondrion 22 (May): 91–95.

Takenaka, Kouhei, Takuma Tanabe, Makoto Kawamukai, and Yasuhiro Matsuo. 2018. “Overexpression of the Transcription Factor Rst2 in Schizosaccharomyces Pombe Indicates Growth Defect, Mitotic Defects, and Microtubule Disorder.” Bioscience, Biotechnology, and Biochemistry 82 (2): 247–57.

Tany, Ryosuke, Yuhei Goto, Yohei Kondo, and Kazuhiro Aoki. 2022. “Quantitative Live-Cell Imaging of GPCR Downstream Signaling Dynamics.” Biochemical Journal 479 (8): 883–900.

Uda, Youichi, Yuhei Goto, Shigekazu Oda, Takayuki Kohchi, Michiyuki Matsuda, and Kazuhiro Aoki. 2017. “Efficient Synthesis of Phycocyanobilin in Mammalian Cells for Optogenetic Control of Cell Signaling.” Proceedings of the National Academy of Sciences of the United States of America 114 (45): 11962–67.

Uda, Youichi, Haruko Miura, Yuhei Goto, Kei Yamamoto, Yusuke Mii, Yohei Kondo, Shinji Takada, and Kazuhiro Aoki. 2020. “Improvement of Phycocyanobilin Synthesis for Genetically Encoded Phytochrome-Based Optogenetics.” *ACS Chemical Biology*, November. 10.1021/acschembio.0c00477.

Uysal Özdemir, Özge, Andrea Krapp, Bastien Mangeat, Marc Spaltenstein, and Viesturs Simanis. 2024. “A Role for the Carbon Source of the Cell and Protein Kinase A in Regulating the S. Pombe Septation Initiation Network.” Journal of Cell Science 137 (1). 10.1242/jcs.261488.

Vandame, Pauline, Corentin Spriet, Dave Trinel, Armance Gelaude, Katia Caillau, Coralie Bompard, Emanuele Biondi, and Jean-François Bodart. 2014. “The Spatio-Temporal Dynamics of PKA Activity Profile during Mitosis and Its Correlation to Chromosome Segregation.” Cell Cycle 13 (20): 3232–40.

Wang, Lili, Kenneth Griffiths, Y. Hi Zhang, F. Douglas Ivey, and Charles S. Hoffman. 2005. “Schizosaccharomyces Pombe Adenylate Cyclase Suppressor Mutations Suggest a Role for cAMP Phosphodiesterase Regulation in Feedback Control of Glucose/cAMP Signaling.” Genetics 171 (4): 1523–33.

Welton, Robert M., and Charles S. Hoffman. 2000. “Glucose Monitoring in Fission Yeast via the gpa2 Gα, the git5 Gβ and the git3 Putative Glucose Receptor.” Genetics 156 (2): 513–21.

Yamawaki-Kataoka, Y., T. Tamaoki, H. R. Choe, H. Tanaka, and T. Kataoka. 1989. “Adenylate Cyclases in Yeast: A Comparison of the Genes from Schizosaccharomyces Pombe and Saccharomyces Cerevisiae.” Proceedings of the National Academy of Sciences of the United States of America 86 (15): 5693–97.

Young, D., M. Riggs, J. Field, A. Vojtek, D. Broek, and M. Wigler. 1989. “The Adenylyl Cyclase Gene from Schizosaccharomyces Pombe.” Proceedings of the National Academy of Sciences of the United States of America 86 (20): 7989–93.

Zhang, Jin, Christopher J. Hupfeld, Susan S. Taylor, Jerrold M. Olefsky, and Roger Y. Tsien. 2005. “Insulin Disrupts Beta-Adrenergic Signalling to Protein Kinase A in Adipocytes.” Nature 437 (7058): 569–73.

Zhang, J., Y. Ma, S. S. Taylor, and R. Y. Tsien. 2001. “Genetically Encoded Reporters of Protein Kinase A Activity Reveal Impact of Substrate Tethering.” Proceedings of the National Academy of Sciences of the United States of America 98 (26): 14997–2.

