## Supplementary Information for "Live-cell fluorescence imaging and optogenetic control of PKA kinase activity in fission yeast *Schizosaccharomyces pombe*"

**Running Title:** Live-cell PKA activity sensor in fission yeast cells

**Authors:**

Keiichiro Sakai^1,2^, Kazuhiro Aoki^1,2,3,4,5§^, Yuhei Goto^1,2,3,§^

**Contact Information:**

^1^Quantitative Biology Research Group, Exploratory Research Center on Life and Living Systems (ExCELLS), National Institutes of Natural Sciences, 5-1 Higashiyama, Myodaiji-cho, Okazaki, Aichi 444-8787, Japan.

^2^Division of Quantitative Biology, National Institute for Basic Biology, National Institutes of Natural Sciences, 5-1 Higashiyama, Myodaiji-cho, Okazaki, Aichi 444-8787, Japan.

^3^Department of Basic Biology, School of Life Science, SOKENDAI (The Graduate University for Advanced Studies), 5-1 Higashiyama, Myodaiji-cho, Okazaki, Aichi 444-8787, Japan.

^4^Center for Living Systems Information Science, Graduate School of Biostudies, Kyoto University, Yoshida-Konoecho, Sakyo-ku, Kyoto, Kyoto 606-8501, Japan

^5^Laboratory of Cell Cycle Regulation, Graduate School of Biostudies, Kyoto University, Yoshida-Konoecho, Sakyo-ku, Kyoto, Kyoto 606-8501, Japan

**Key words:** fission yeast, cAMP-PKA pathway, kinase translocation reporter (KTR), optogenetics

### **Supplementary movies**

### **Movie S1. Translocation of the KTR-based PKA biosensor, spPKA-KTR1.0, in response to glucose stimulation in fission yeast cells.**

Representative confocal fluorescence images of spPKA-KTR1.0 (left) and Hta1 (histone H2A) fused to mNeonGreen (Hta1-mNG, right) in fission yeast cells. Cells were cultured in YEA supplemented with 2% glycerol, and then stimulated with YEA containing 2% glucose from 30 minutes to 60 minutes (marked with “+ Glucose” in the movie). Fluorescence images were taken every one minute, and the playback speed was set to 40 frames per second. The time stamp indicates the time in hours: minutes. Scale bar, 10 μm.

### **Movie S2. Optogenetic control of PKA kinase activity by using a photoactivatable adenylate cyclase bPAC in fission yeast cells.**

Representative confocal fluorescence images of the *cyr1*Δ strain expressing the spPKA-KTR1.0 (left), NLS-iRFP-NLS (right), and bPAC. Cells were cultured under dark conditions in 2% glucose-supplemented YEA medium and then stimulated with blue light (labeled “Blue light” in the movie). Fluorescence images were taken every 30 seconds, and the playback speed is set to 40 frames per second. The time stamp indicates the time in hours: minutes. Scale bar, 10 μm.

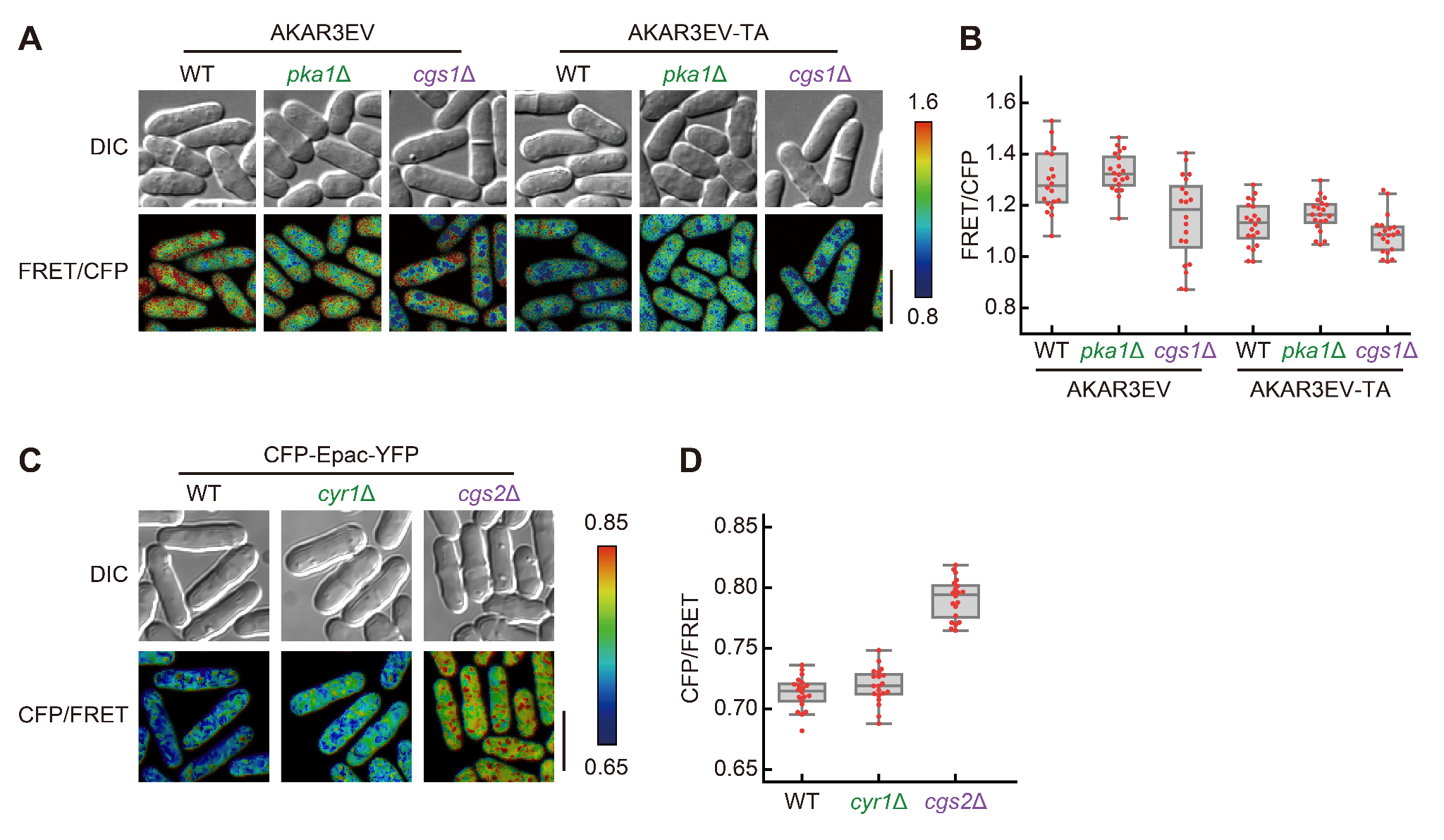

**Supplementary Figure 1. Characterization of the FRET-based PKA biosensor AKAR3EV and the cAMP biosensor CFP-Epac-YFP in *S. pombe*.**

(A) AKAR3EV and a non-phosphorylatable mutant AKAR3EV-TA in wild-type, *pka1*Δ, and *cgs1*Δ cells. AKAR3EV is a FRET-based PKA biosensor, which was originally developed to measure the PKA activity in mammalian cells (Komatsu et al. 2011). AKAR3EV has a substrate domain that is phosphorylated by PKA (Jin Zhang et al. 2005). AKAR3EV is phosphorylated by PKA, undergoes conformational changes, and exhibits a higher FRET/CFP ratio than that in the non-phosphorylated state. In AKAR3EV-TA, a PKA phosphorylation site (T506) is substituted with an alanine residue, and therefore the FRET/CFP ratio is constantly low regardless of the PKA activity. Representative FRET/CFP ratio images of AKAR3EV and AKAR3EV-TA are shown in the intensity-modulated display mode with DIC images. Scale bar, 10 μm. (B) FRET/CFP ratios of AKAR3EV and AKAR3EV-TA were quantified from 20 cells. Each dot represents the FRET/CFP ratio of a single cell. The FRET/CFP of AKAR3EV-TA was lower than that of wild-type AKAR3EV in the rich medium, indicating that the phosphorylation of AKAR3EV increases the FRET/CFP in fission yeast cells cultured in rich medium. However, the FRET/CFP of AKAR3EV was not affected by the deletion of either *pka1* or *cgs1*, genes encoding the catalytic and regulatory subunits of PKA, respectively. These results suggest that AKAR3EV is not phosphorylated by PKA but by the other kinases. (C) CFP-Epac-YFP in wild-type, *cyr1*Δ, and *cgs2*Δ cells. CFP-Epac-YFP is FRET-based cAMP biosensor (Ponsioen et al. 2004), and shows reduced FRET efficiency in response to cAMP elevation. Representative CFP/FRET ratio images are shown in the intensity-modulated display mode with DIC images. Scale bar, 10 μm. (D) CFP/FRET ratios of CFP-Epac-YFP were quantified from 20 cells. Each dot represents the CFP/FRET ratio of a single cell. CFP-Epac-YFP in fission yeast grown in rich medium did not significantly decrease in the CFP/FRET ratio even in *cyr1*Δ cells, in which cAMP levels are low due to the absence of adenylate cyclase. These data suggest that the basal cAMP levels in fission yeast cells are lower than the sensitivity of the sensor. In contrast, CFP/FRET was significantly increased by the deletion of a phosphodiesterase gene *cgs2*, indicating that CFP-Epac-YFP detects higher cAMP levels rather than basal and low cAMP levels in WT and *cyr1*Δ cells, respectively. Therefore, CFP-Epac-YFP cannot be used to measure the physiological dynamics of cAMP levels in fission yeast cells.

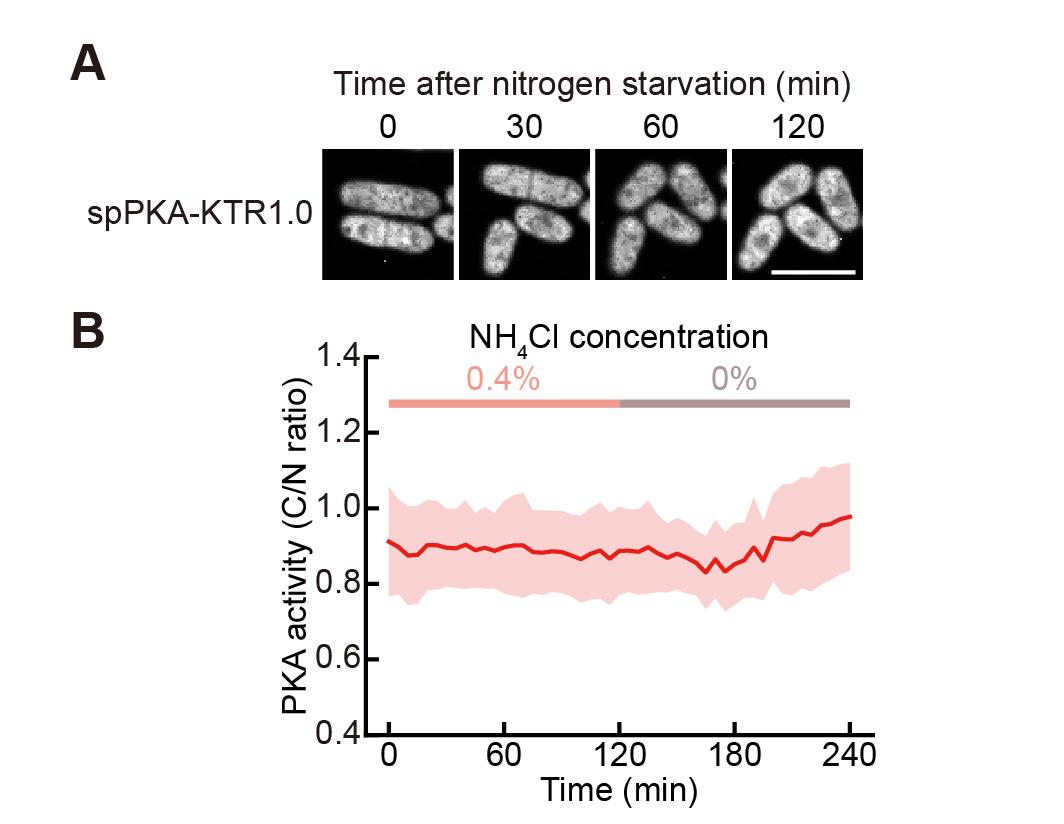

**Supplementary Figure 2. Nitrogen starvation did not affect the PKA dynamics in fission yeast cells.**

(A) Representative confocal fluorescence images of the spPKA-KTR1.0 during nitrogen starvation. Cells were cultured in EMM containing 2% glucose and 0.4% NH_4_Cl, and then medium was changed to EMM containing 2% glucose and no NH_4_Cl. Scale bar, 10 μm. (B) The C/N ratio of spPKA-KTR1.0 (n = 25 cells) during nitrogen starvation was calculated, and the mean values of C/N ratios are shown with error bars (SD).

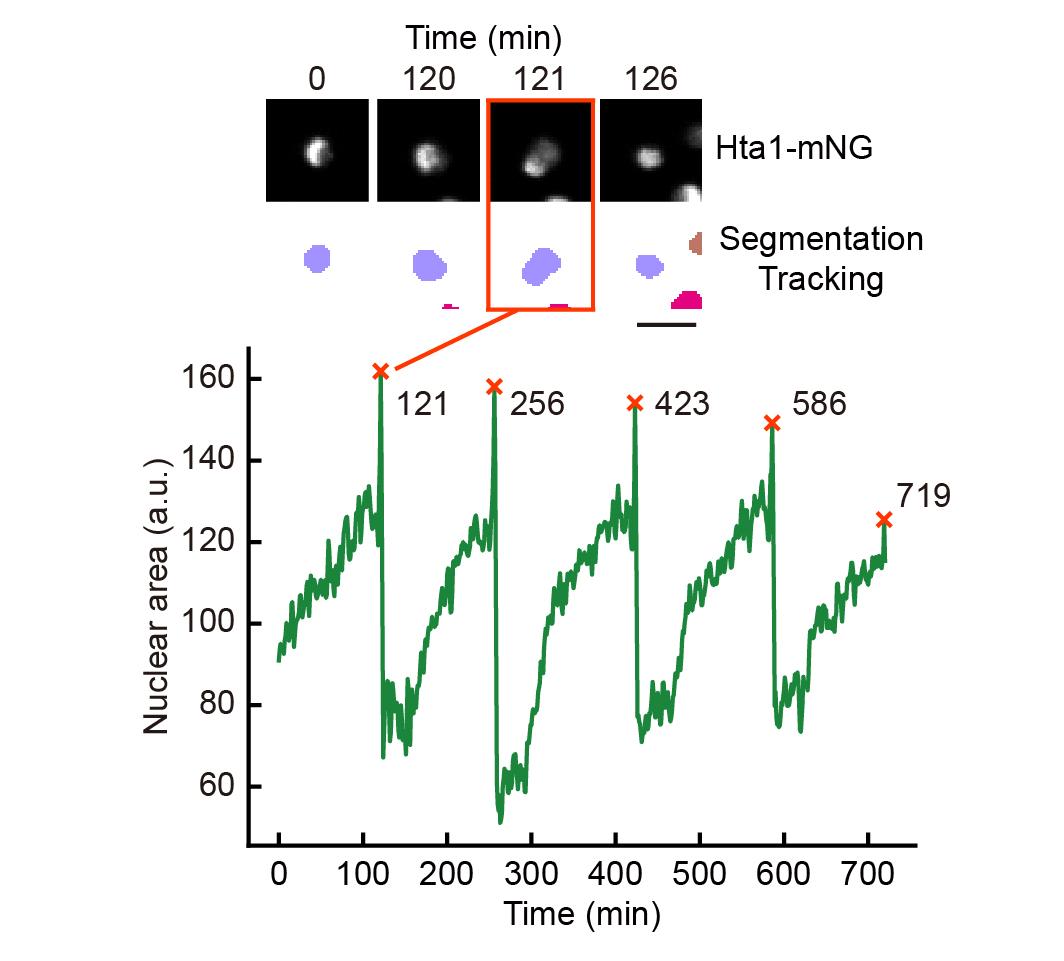

**Supplementary Figure 3. Segmentation, tracking, and cell division timing detection using the nuclear marker.**

Representative nuclear images (Hta1-mNG) and segmentation of nuclear region (top). At the time just before the nuclear division (for example, 121 min), the nuclear area became expanded. The quantification of the nuclear area shows clear peaks at the timing of nuclear division (bottom).

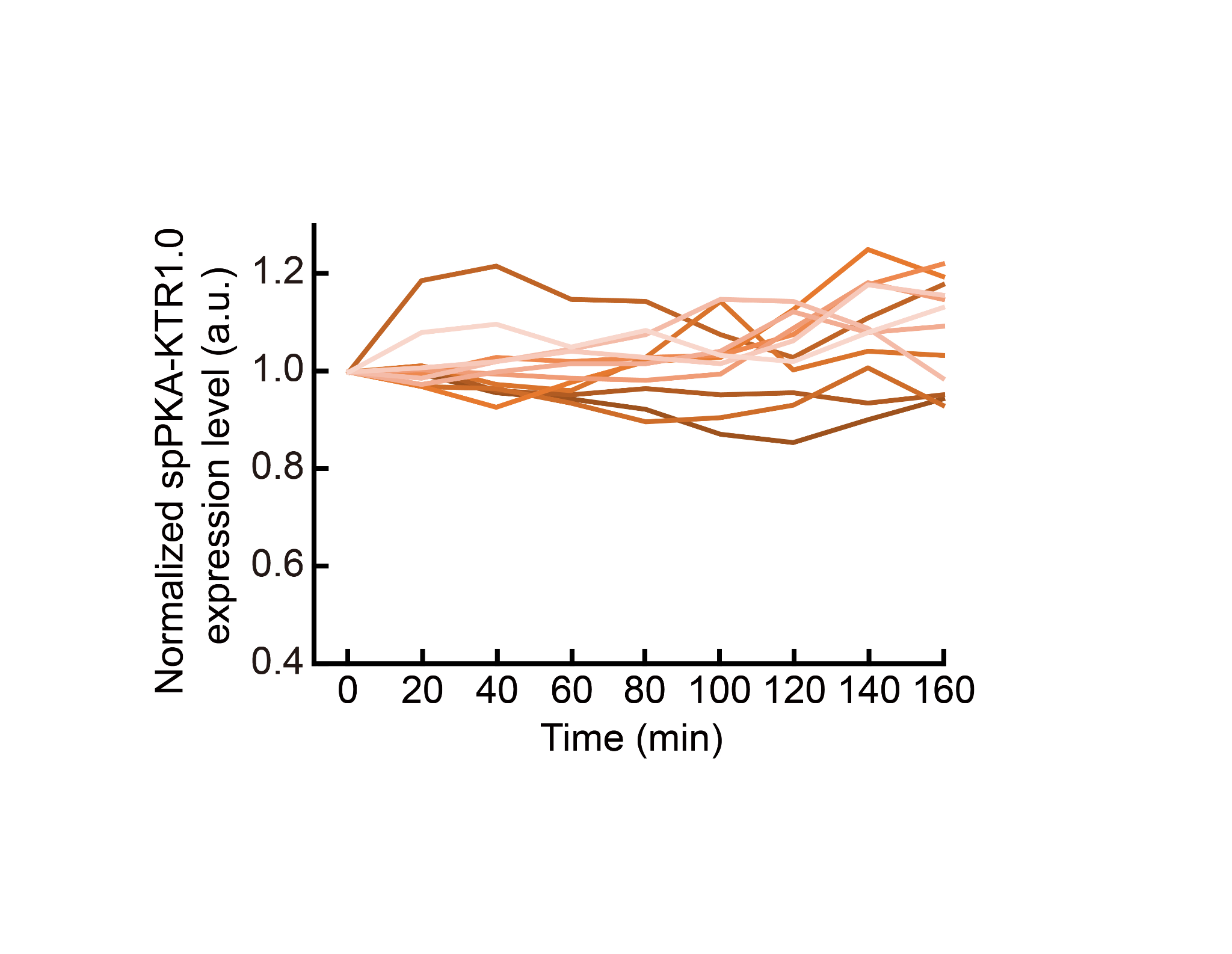

**Supplementary Figure 4. Expression level of spPKA-KTR1.0 throughout the cell cycle.**

Signal intensities of spPKA-KTR1.0 for 160 minutes were quantified from 12 cells. Fluorescence intensities were normalized by the intensity at time zero.
